# CD155 and EndoA1 mediate growth and tissue invasion downstream of MAP4K4 in medulloblastoma cells

**DOI:** 10.1101/2021.08.10.455785

**Authors:** Charles Capdeville, Linda Russo, David Penton, Jessica Migliavacca, Milica Zecevic, Alexandre Gries, Stephan C.F. Neuhauss, Michael A. Grotzer, Martin Baumgartner

**Affiliations:** Pediatric Molecular Neuro-Oncology Lab, Children’s Research Center, University Children’s Hospital Zürich, Zürich, Switzerland; Department of Molecular Life Sciences, University of Zurich, Zürich, Switzerland; Department of Oncology, University Children’s Hospital Zürich, Zürich, Switzerland

**Keywords:** CD155/PVR, Endophilin A1, MAP4K4, Medulloblastoma, Tissue invasion

## Abstract

The composition of the plasma membrane (PM)-associated proteome of tumor cells determines cell-cell and cell-matrix interactions and the response to environmental cues. Whether the PM-associated proteome impacts the phenotype of Medulloblastoma (MB) tumor cells and how it adapts in response to growth factor cues is poorly understood. Using a spatial proteomics approach, we observed that hepatocyte growth factor (HGF)-induced activation of the receptor tyrosine kinase c-MET in MB cells changes the abundance of transmembrane and membrane-associated proteins. The depletion of MAP4K4, a pro-migratory effector kinase downstream of c-MET, leads to a specific decrease of the adhesion and immunomodulatory receptor CD155 and of components of the fast-endophilin-mediated endocytosis (FEME) machinery in the PM-associated proteome of HGF-activated MB cells. The decreased surface expression of CD155 or of the FEME effector Endophilin A1 reduces growth and invasiveness of MB tumor cells in the tissue context. These data thus describe a novel function of MAP4K4 in the control of the PM-associated proteome of tumor cells and identified two downstream effector mechanisms controlling proliferation and invasiveness of MB cells.

**Graphical abstract:** c-MET activation upon HGF stimulation induces c-MET internalization and induces downstream MAP4K4 activity. **(1)** MAP4K4 is required downstream of activated c-MET for the maintenance of surface presentation of CD155 in activated cells. CD155 expression is required for MB cell migration, invasion and proliferation in the tissue context. **(2)** MAP4K4 is required downstream of activated c-MET to maintain membrane depolarization, possibly by regulating the surface localization of several ion channels and transporters. **(3)** MAP4K4 is required downstream of activated c-MET cause PM-proximal localization of FEME effector CIP4, FBP17 and CIN85. The FEME effector endophilin A is necessary for MB cell migration, invasion and dissemination.

## Introduction

The plasma membrane (PM)-associated proteome is a dynamic interface of mammalian cells, which orchestrates interactions with the extracellular environment and transmits extracellular cues. This physical barrier regulates the transfer of chemical and mechanical information between extracellular and intracellular spaces. The net outcome of the functions of PM-associated proteins affects the cellular phenotype and the composition and biophysical structure of the extracellular microenvironment. The PM-associated proteome primarily consists of receptors, cell adhesion molecules, enzymes and ion channels^1^, and their expression defines the strength and duration of signal transmission and how the cell interacts with its environment. Besides these transmembrane proteins, numerous proteins are associated with the PM either directly through lipid anchors, or indirectly by binding to transmembrane proteins. The diversity of proteins associated with the PM and their differential expression, localization, or regulation in pathological conditions such as cancer not only constitutes the condition-specific biomarker landscape^2^, but also defines a druggable repertoire of therapeutic targets to interfere with the oncogenic phenotype.

The PM-associated proteome of a cell is dynamically controlled through membrane turnover via endocytic uptake and exocytic fusion^3, 4^, which involves the dynamic reorganization of the cortical cytoskeleton. Membrane trafficking and cytoskeleton reorganization are thus key processes that position chemical and mechanical sensors and enable the uptake of nutrients or extrusion of signal transmitters, thereby determining the cellular response to complex environments. Moreover, endocytosis and the sequential intracellular vesicular trafficking control adhesion turnover, signal transduction, consequently influencing the duration of mitogenic or mobile phenotypes. De-regulation of physiological trafficking in tumor cells may thus contribute to cell transformation^5^, and especially altered endocytic activity can broadly impact tumor cell growth, invasion and metastasis, and viability^6, 7^.

Medulloblastoma (MB) is a malignant embryonal neuroepithelial tumor of the cerebellum with a high propensity to disseminate and metastasize within the central nervous system^8^. Several omics studies described the genomic heterogeneity in MB, which is now classified into four subgroups with twelve subtypes in total^9, 10^. Despite this molecular understanding of MB, the current standard therapy still consists of surgical resection of the tumor and craniospinal irradiation followed by chemotherapy^11^. These anti-cancer therapies can have a devastating impact on the developing brain of the patients and can cause severe long-term side effects such as cognitive or mobility impairments^12^. Thus, novel therapeutic options with phenotype-specific anti-tumor activity are needed to replace or complement current treatments and reduce therapy-associated side effects.

The Ser/Thr kinase MAP4K4 is overexpressed in MB and contributes to the pro-invasive phenotype of MB cells downstream of hepatocyte growth factor (HGF)-cellular mesenchymal-epithelial transition factor (c-MET) stimulation^13, 14^. c-MET receptor tyrosine kinase (RTK) activation caused increased membrane dynamics, increased endocytic uptake and the activation and turnover of integrin ß1 adhesion receptor^13^. Integrin activation and internalization of integrins are necessary for cell migration and invasion^15, 16^. Internalization of activated RTKs and integrins furthermore modulates pathway activation at endosomal compartments^17, 18^. Thereby, the condition-specific endocytic activity adds a layer of regulatory impact on tumor cell behavior.

Endocytic cellular processes like synaptic vesicle endocytosis, receptor trafficking, apoptosis and mitochondrial network dynamics can be controlled by endophilins, which are members of the BAR (Bin/amphiphysin/Rvs) domain superfamily ^19, 20^. Dysregulation of these processes is associated with cancer or neurodegenerative diseases^19, 21^. Endophilins were initially considered a peripheral component of clathrin-mediated endocytosis^22, 23^. A novel clathrin-independent endocytic pathway based on endophilin A subfamily activity, termed fast endophilin mediated endocytosis (FEME) was described recently^21^, and several studies also implicated endophilins in cell motility control^24–26^. The endophilin A1 is almost exclusively expressed in brain tissue^27^

How the PM proteome of MB cells is regulated in response to RTK activation, and which potential therapeutic targets may be represented in the dynamic PM compartment are unknown. Therefore, we used a spatial proteomics approach^28^ and determined the PM-associated proteome of resting and RTK-activated MB cells. We addressed whether the pro-oncogenic activity of MAP4K4 could be coupled to RTK-induced, dynamic alterations in the PM proteome. This analysis allowed the identification of PM-associated proteins subjected to c-MET-induced spatial re-distribution and the identification of MAP4K4 as a regulator of this process. We describe MAP4K4-dependent alterations in PM association of solute carriers, ion channels and of the immune modulatory receptors. Furthermore, we address underlying mechanisms and potential functional consequences of this regulation for cancer cell homeostasis and cancer cell motility.

## Results

### HGF-c-MET signaling promotes PM proteome reorganization in medulloblastoma cells

To determine changes in PM and PM-proximal association of proteins in response to growth factor stimulation, we used a PM-associated proteome analysis approach (Fig. 1A). We used membrane-impermeable Sulfo-NHS-SS-biotin labeling of intact sgControl (sgCTL, sgC) and sgMAP4K4 (MAP4K4 knockout, sgM4, Fig S1A,B) MB tumor cells, followed by streptavidin protein capture and high-resolution mass spectrometry analysis (Fig. 1A). The data generated a rather diverse set of proteins, which is available via ProteomeXchange with the identifier PXD030597. We observed no significant difference in the PM proteome between sgCTL and sgMAP4K4 cells under starved conditions (Fig. S1C). Stimulation with HGF caused an alteration in protein abundance (Fig. S1C,D), suggesting that growth factor stimulation alters the composition of PM-associated proteins.. To specifically enrich for membrane or membrane-associated proteins, we filtered the data either by using the transmembrane prediction algorithm TMHMM^29^ or by annotating the proteins according to their known subcellular localization^30^. These stringent sorting criteria identified 952 TM proteins and 336 PM-annotated proteins, respectively (Fig. 1A, Tables 1 & 2). In sgCTL cells, HGF stimulation decreased the abundance of a number of TM (Fig. 1B, S1D, Table 3) or PM-annotated proteins (Fig. 1B, S1D, Table 4). This indicated that either internalization of these proteins in response to c-MET activation is increased or recycling decreased. We previously found that internalization of activated c-MET depends on MAP4K4^13^. Therefore, we tested whether MAP4K4 could contribute to reduced PM association in cells, where c-MET is activated by HGF stimulation. By comparing HGF-stimulated sgCTL and sgMAP4K4 cells, we detected changes in the abundance of >1500 proteins (fold change >2, adjusted p-value <0.2, Fig. 1C, S1F). Specifically, HGF stimulation in MAP4K4 depleted cells led to an increased abundance of TM proteins (Fig. 1B,C, Table 3) and PM-annotated proteins (Fig. 1B,C, Table 4), suggesting a regulatory function of MAP4K4 in PM localization of these proteins. A paradigmatic example is c-MET itself, which is internalized in a MAP4K4-dependent manner in DAOY cells^14^ and detected at lower levels in the PM proteome in response to HGF stimulation in sgCTL cells (Fig. 1D). MAP4K4 contributes to this process as depletion of MAP4K4 significantly increases c-MET abundance on the PM in HGF-stimulated cells. None of the five additional RTKs detected by mass-spectrometry displayed similar behavior (Fig. 1D). We concluded from this experiment that c-MET activation triggers the turnover of PM-associated proteins and that MAP4K4 is indispensable for the turnover of a number of those proteins in MB cells. Gene ontology analysis of PM-associated proteins revealed MAP4K4 dependencies in several pathways controlling endo- and exocytotic activities of cells as well as immune cell recognition or solute exchange (Fig. 1E). It also revealed that HGF stimulation in sgMAP4K4 cells increased PM association of proteins with reported implications in chemotherapy sensitivity/resistance (SLC29A1, SLC4A7, LRRC8)^31, 32^ (Fig. 1F, Table 3). Conversely, MAP4K4 depletion decreased PM association of proteins involved in tumor immune evasion (CD155, CD276, HLA-G)^33–35^, in solute influx regulation (CLIC1, SLCs, ABCCs)^36, 37^ and cell migration (CD155, CD276)^33, 38^ (Fig. 1F, tables 1-4). In conclusion, we found that MET activation by HGF triggers reorganization of the PM-associated proteome in MB cells and that MAP4K4 is involved in the selective control of this process.

**Figure 1.**
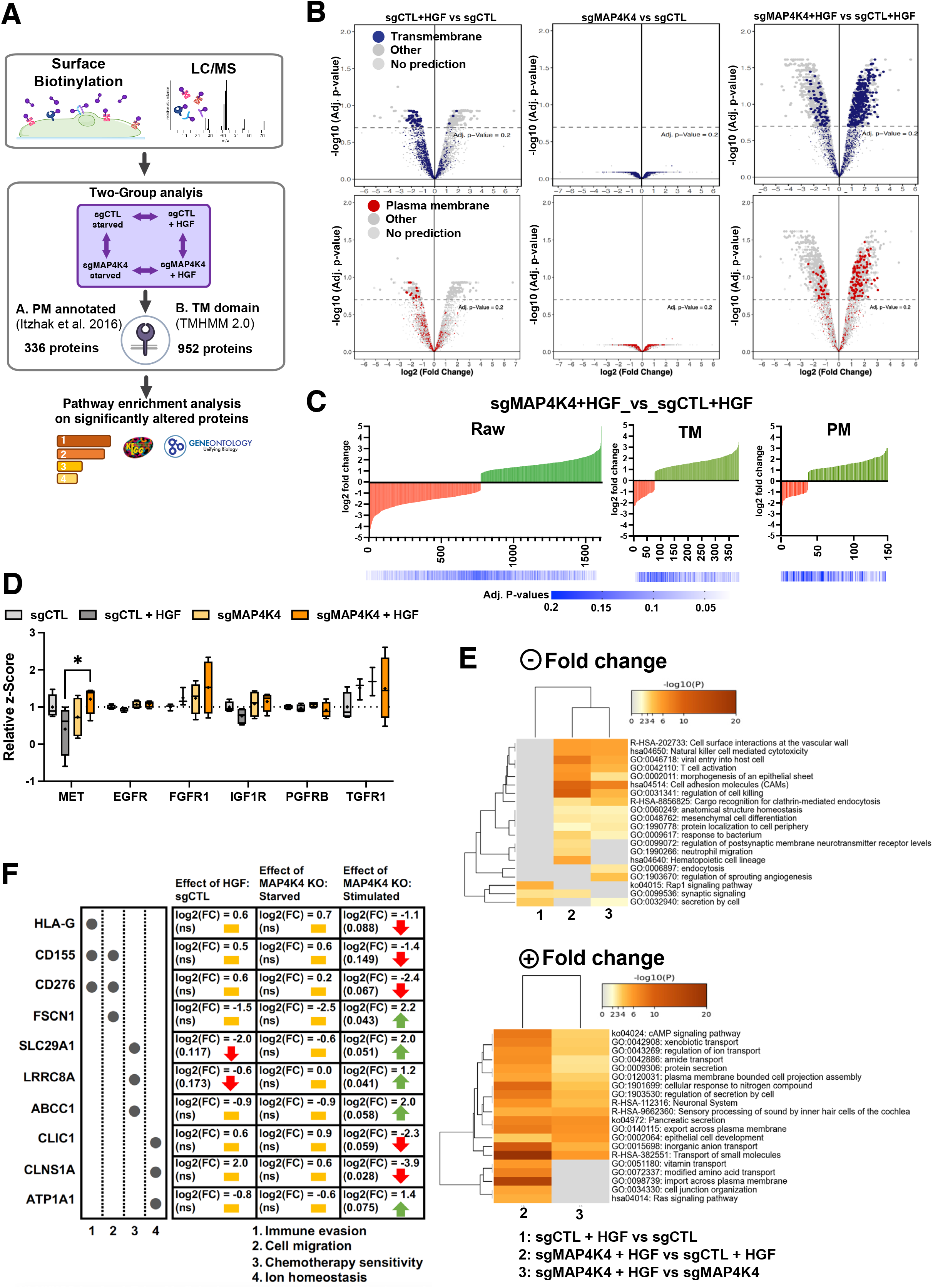
HGF-c-MET signaling alters composition of plasma membrane (PM)-associated proteome. **(A)** General workflow for cell-surface protein isolation, mass spectrometry and computational analysis used. **(B)** Volcano plot of PM-associated proteins identified by deep mass-spectrometry analysis. Colored dots indicate proteins with predicted TM domain (blue, upper) or PM localization (red, lower). sgCTL: CRISPR/CAS-9 control guide RNA expressing cells; sgMAP4K4: CRISPR/CAS-9 MAP4K4-specific guide RNA expressing cells; + HGF: SFM supplemented with 20 ng/ml HGF for 30 min. **(C)** log2 Fold change of RAW, TM or PM protein comparisons between treatments. Green: positive fold change (increased association), red: negative fold change (decreased association). **(D)** Quantification of the PM association of different receptor tyrosine kinases. Box plots display normalized z-score values extracted from all-group comparison of MS analysis. N = 4 independent experiments, min to max, * = p<0.05 (mixed effect analysis, with Šídák’s multiple comparisons test). **E)** Heatmaps of enriched ontology clusters across multiple two-group analyses of significantly altered, PM-annotated proteins with positive (upper) or negative (lower) fold change. Panel colors are defined by adjusted p-values. Gray cells: No enrichment detected for that term in the corresponding two-group analyses. Missing comparisons: no enrichment detected **(F)** Overview of a selection of potentially clinically relevant PM-associated proteins (log2 FC >1, adjusted p-values <0.2). Yellow bar: no change; red arrow: decreased PM association; green arrow: increased PM association.

### Altered transporter expression in MAP4K4-depleted cells does not significantly affect sensitivity to Etoposide and Lomustine

To better understand which protein functions are particularly affected by the altered PM-associated proteome, we annotated the filtered proteins using Protein Atlas^39, 40^. We found that 223 of 952 transmembrane proteins (Fig. S2A) and 84 of 336 plasma membrane proteins (Fig. S2B) are annotated as transporters. These trans-membrane proteins mediate the transfer of ions, small molecules, and macromolecules across membranes^41^, thus suggesting the potential implication of the HGF-c-MET-MAP4K4 axis in maintaining ion homeostasis of MB cells, and in transferring nutrients and drugs across the PM. Dysregulation of influx or efflux mechanisms can lead to chemotherapy sensitivity or resistance in cancer cells^42, 43^. We detected altered PM association of transporters including SLC29A1, SLC4A7 and LRRC8A, where contribution to the influx/efflux of standard chemotherapy cytotoxic drugs has been described^31–32, 44–45^. Depletion of MAP4K4 prevented the reduced PM-association of these transporters after HGF stimulation (Fig. S2C). The differences in PM association are not the consequences of altered regulation of gene expression (Fig. S1B). We hence tested the susceptibilities of the cells to cytotoxic drugs by comparing dose-response curves for Lomustine^46^ and Etoposide^47^ of sgCTL and sgMAP4K4 cells. Under normal growth conditions (10% FBS), depletion of MAP4K4 did not significantly impact the sensitivity of DAOY cells to Etoposide (IC50 value of 3.2 µM and 2.8 µM respectively) or Lomustine (IC50 value of 46.2 µM and 39.5 µM respectively) (Fig. S2D). We additionally tested drug response in cells maintained first in low serum media for 24h (1% FBS) and then treated the cells for 48h with HGF in combination with Lomustine or Etoposide. Under these conditions, depletion of MAP4K4 caused some reduction in Lomustine sensitivity (Fig. S2E). We observed no impact on the response to Etoposide treatment. Taken together, these results suggest a weak sensitizing effect of MAP4K4 towards Lomustine.

### MAP4K4 controls PM-association of ion transport proteins and prevents cell hyperpolarization in HGF-stimulated cells

We furthermore detected MAP4K4-dependent PM association of the chloride channel CLIC1 in HGF-stimulated cells (Fig. 2A, S2F). CLIC1 contributes to MB proliferation *in vitro*, and CLIC1 depletion in MB tumor cell lines significantly reduces tumor cell growth *in vivo*^37^. Gene expression analysis of several MB patient sample cohorts shows CLIC1 overexpression in the tumor samples compared to control tissues (Fig. 2B). Importantly, this elevated CLIC1 expression is associated with worse outcomes in SHH-alpha and Gr4 MB tumors (Fig. 2C). In control cells, PM association of CLIC1 remains unaltered by HGF stimulation. In contrast, HGF stimulation of MAP4K4-depleted cells caused a significant drop of CLIC1 PM association (Fig 2A), suggesting that MAP4K4 is either necessary for CLIC1 transfer to or maintenance at the PM in growth factor-stimulated cells. Conversely, the PM-association of the acetyl-CoA transporter SLC33A1, the H^+^/Ca2^+^/Mn2^+^ exchanger TMEM165, the H^+^/K^+^ transporter ATPase ATP12A, the sodium bicarbonate co-transporter SLC4A7, the acetyl-CoA transporter SLC19A1 and the Na+/K+ transporter ATP1A3 is significantly increased in MAP4K4-depleted cells stimulated with HGF (Fig. 2D). Using RT-qPCR we found no evidence that MAP4K4 depletion altered the mRNA expression of a selection of proteins identified in our study, indicating that altered PM-association is not the consequence of altered transcriptional control (Fig. S1B).

**Figure 2.**
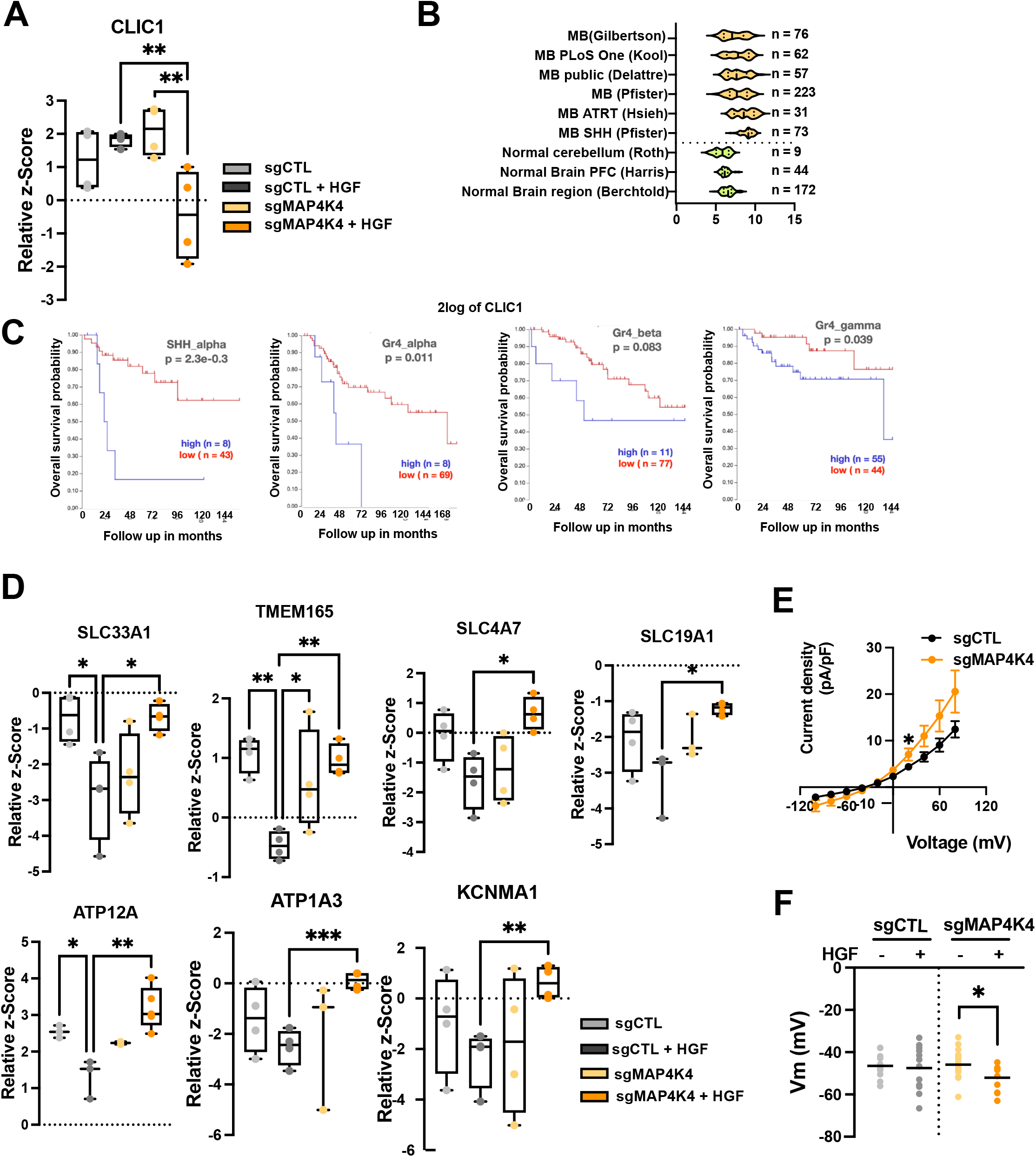
MAP4K4 modulates the ion homeostasis of DAOY cells. **(A)** Quantification of PM association of CLIC1 by MS. Box plots display normalized z-score values extracted from all-group comparison of MS analysis. N = 4 independent experiments, min to max, ** = adjusted p<0.01 of one-way ANOVA. **(B)** U133P Affymetrix gene chip micro array analysis of CLIC1 mRNA expression levels in normal brain tissue and MB tumor samples. **(C)** Kaplan Meier survival analyses in four MB subgroups relative to CLIC1 mRNA expression levels. **(D)** Quantification of PM association of ion transport proteins by MS. Box plots display normalized z-score values extracted from all-group comparison of MS analysis. N = 4 independent experiment, min to max, * = adjusted p<0.05, ** = adjusted p<0.01 of one-way ANOVA. **(E)** Whole cell currents measured at holding potential ranging from −100 mV to +80 mV (in steps of 20 mV) in DAOY cells +/- KO of MAP4K4. N = 13-16 cells measured independently. **F)** Quantification of the membrane potential of DAOY cells +/- KO of MAP4K4 and +/- HGF stimulation (20 ng/ml, 30 min) in normal [Cl^-^] SFM medium. N = 8-16 cells measured independently.

To investigate the significance of this HGF- and MAP4K4-dependent modulation of PM localization of ion channels in MB cell ion homeostasis, we performed whole-cell patch-clamp measurement of the electrical properties of sgCTL and sgMAP4K4 DAOY cells. Under starved conditions, MAP4K4 depletion led to an increased whole-cell current compared to control cells (Fig. 2E). As we observed the most striking differences in PM association between sgCTL and sgMAP4K4 cells after HGF stimulation, we quantified the resting membrane potential (V_mem_) under these conditions. We found that HGF stimulation caused hyperpolarization of MAP4K4-depleted compared to control cells (Fig. 2F). This last finding is in accordance with a more depolarized V_mem_ in disseminating cancer cells, where hyperpolarization leads to a loss of metastatic potential^48^.

Taken together, these data indicated the implication of MAP4K4 in modulating PM association of ion transport proteins in growth factor-stimulated cells and that one consequence of this MAP4K4 function is to prevent hyperpolarization after HGF stimulation.

### Reduced PM association of CD155 correlates with reduced invasion and motility of MB cells

We found differential PM association of the adhesion and immune-modulatory proteins CD155 and CD276 (Fig. 1F). CD276 impacts colorectal cancer cell migration through the Jak2/Stat3/MMP-9 signaling pathway^38^. CD155, also referred to as poliovirus receptor (PVR), is a member of the nectin-like family of proteins that can interact with integrins or RTKs and increases FAK, Ras, or RAP1 downstream signaling^49^. CD155 is involved in cell adhesion, migration, proliferation, or immune regulation^33^, and it is often upregulated in tumors compared to healthy tissues^49^. This renders CD155 particularly interesting in the context of tumor progression, where it could contribute to immune evasion and dissemination.

HGF stimulation in sgMAP4K4 cells causes reduced PM abundance of CD155 compared to sgCTL cells (Fig. 3A). Decreased PM association of CD155 in MAP4K4 depleted cells is not the result of transcriptional control of *CD155* (Fig. 3B) or reduced protein expression (S1E,H). HGF stimulation rather increased mRNA expression of CD155 both in sgCTL and sgMAP4K4 cells by 10 – 20% (Fig. 3B), excluding the possibility that the decrease in CD155 PM association in HGF-stimulated sgMAP4K4 is a consequence of transcriptional repression. Combined these findings indicated that MAP4K4 is either required for the transfer or the maintenance of CD155 at the PM under conditions that trigger membrane dynamics and endocytic turnover.

**Figure 3.**
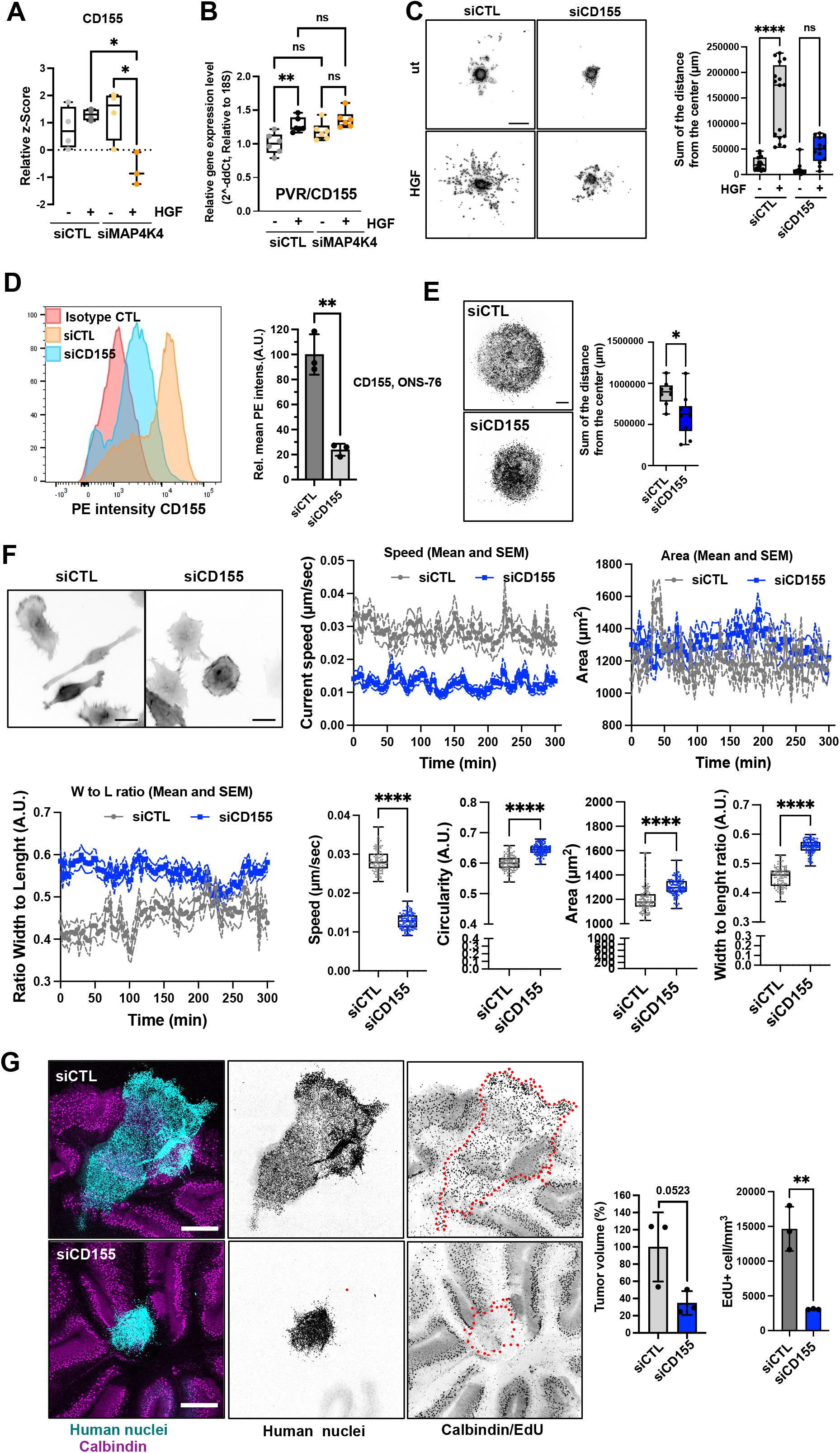
CD155 promotes migration and invasion. **(A)** Quantification of PM association of CD155. Box plots display normalized z-score values extracted from all-group comparison of MS analysis. N = 4 independent experiments, min to max, * = p<0.05 of one-way ANOVA. **(B)** RT-qPCR analysis of *CD155* mRNA expression in siCTL and siMAP4K4 -/+ HGF stimulation (30 min), ** = p<0.01 of one-way ANOVA. **(C)** SIA of siCTL or siCD155 transfected DAOY cells -/+ HGF stimulation. Left: Representative images of Hoechst-stained nuclei in inverted greyscale at end point of SIA. Scale bar = 300 µm. Right Box plots display sum of distances of invasion. N = 3 independent experiments, min to max, * = p<0.05, **** = p<0.0001 of one-way ANOVA. **(D)** Surface expression analysis of CD155 by flow cytometry on ONS-76 cells. Left: Histograms of anti-CD155 PE intensities. Right: bar diagram depicting mean PE (CD155) intensities in siCD155 relative to siCTL cells of n = 3 independent experiments, ** = p<0.01 of unpaired t-test. **(E)** SIA of siCTL or siCD155 transfected ONS-76 cells. Left: Representative images of Hoechst-stained nuclei in inverted greyscale. Scale bar = 300 µm. Right Box plots display sum of distances of invasion. N = 3 independent experiments. * = p<0.05 of unpaired t-test. **(F)** Single cell motility analysis of siCTL and siCD155 transfected ONS-76 cells. Inverted grey-scale images are representative LA-EGFP fluorescence images of siRNA transfected DAOY cells seeded on collagen-I coated plates. Scale bar = 25 µm. Speed and morphological properties of cells were quantified in 5 min intervals for 300 min and are displayed over time (x/y plots) and averaged (box-dot plots). N = 3 independent experiments, min to max, **** = p<0.0001 of unpaired t-test, ns = not significant. **(G)** OCSC of siCTL or siCD155 ONS-76 cells implanted on cerebellar slices for 48h. Left: representative images of OCSCs. Cyan: Human nuclei, magenta: Calbindin (Purkinje cells), Scale bar: 400 µm. Right: Bar plots show volume and proliferation index of n = 3 spheroids ** = p<0.01 (unpaired t-test).

To test the requirement of CD155 for HGF-induced collagen invasion, we performed a spheroid invasion assay (SIA)^50, 51^ (Fig. 3C) using CD155-depleted DAOY cells (Fig. S1H). We found that the reduction of CD155 expression stalled HGF-induced collagen I invasion. To corroborate the implication of CD155 in migration control with another cell line, we tested the effect of CD155 depletion on collagen I invasion in ONS-76 SHH MB cells. These cells invade collagen I independently of exogenous growth factor stimulation. We first confirmed CD155 depletion by siRNA using FACS (Fig. 3D). We then found that CD155 depletion significantly reduced collagen I invasion (Fig. 3E) and that it also reduced proliferation of ONS-76 cells grown as spheroids in 3D cultures (Fig. S3A) without inducing cell death (Fig. S3B). As the inhibition of proliferation by Mitomycin C did not block collagen I invasion in ONS-76 cells (Fig. S3C), we concluded that the reduced disseminated cell count in the collagen invasion analysis of CD155-depleted cells is not primarily a consequence of reduced proliferation, but rather of a reduced migratory potential of these cells. To further test this possibility, we assessed the consequence of reduced CD155 expression on motility parameters more directly by single-cell analysis of CD155-depleted ONS-76 cells seeded on collagen I coated plates. Cell movements were recorded by time-lapse video microscopy for five hours. CD155 depletion reduced the average speed of ONS-76 cells by 50% (Fig 3F). Phenotypically, cells with decreased CD155 expression displayed increased circularity, increased area, and increased width to length ratio (Fig 3F). Taken together, these data identified the CD155 cell adhesion molecule as a promotor of proliferation, motility, and invasiveness in SHH MB cell models.

### CD155 is necessary for cerebellar tissue infiltration

To assess whether CD155 depletion prevented cerebellar tissue infiltration and growth of MB cells, we implanted ONS-76-LA-EGFP tumor spheroids onto cerebellar slices and determined growth and invasion after tumor-cerebellar slice co-culture^52^. After 48 hours of co-culture, ONS-76-LA-EGFP cells transfected with siCTL displayed sheet-like infiltration of the surrounding brain tissue (Fig. 3G). In contrast, spheroids of cells transfected with siCD155 did display a much-reduced invasive behavior and remained clustered at the site of initial spheroid positioning. As an indirect measurand for tumor growth and invasion, we quantified the volume of the tumor cell clusters. We observed a ∼2-fold reduction of the cluster volume in siCD155 transfected compared to siCTL cells (Fig. 3G, S3D). Some cells in the siCD155 condition still display an invasive phenotype (Fig, 3G, S3D), possibly due to variations in transfection efficiency at the single-cell level. We also evaluated tumor cell proliferation in the tissue-embedded tumor cell clusters using EdU incorporation. We found that depletion of CD155 caused a significant decrease in the number of proliferative cells in the *ex vivo* model, indicating that CD155 also contributes to proliferation in the tissue context (Fig. 3G, S3D). Together, these findings demonstrate that the cell adhesion receptor CD155 contributes to tissue invasion and growth in SHH MB cells.

### Increased expression of MAP4K4 and EndoA1 in SHH MB

MAP4K4 is overexpressed in MB patient tumor samples^14^ and gene expression profiling of 763 primary MB samples^53^ revealed that *MAP4K4* expression correlates positively with genes involved in endocytosis control (Fig. S4A). Interestingly, the strongest positive correlate with MAP4K4 expression among the endocytosis-regulating genes is *SH3GL2* (EndoA1, r = 0.501), while expression of both *CLTB* (r = - 0.228) and *CLTC* (r = - 0.228) correlate negatively with *MAP4K4* (Fig. S4B). The highest positive correlation between *SH3GL2* and *MAP4K4* expression is observed in SHH MB tumors, where also *CLTC* and *SH3GL2* expression correlate negatively (r=-0.144) (Fig. 4A). Thus, decreased cell migration and altered surface proteome composition in MAP4K4-depleted cells could be mechanistically linked to a clathrin-independent mechanism of endocytosis downstream of HGF-c-MET signaling. A recent genomic and proteomic analysis of a cohort of 218 pediatric brain tumors^54^ revealed increased levels of MAP4K4, EndoA1, and EndoA3 – both at protein and RNA levels – but not of EndoA2, in primary MB and ganglioglioma tumor samples (Fig. S4C). These findings highlight high expression of endophilins in MB compared to other central nervous system tumors and pointed towards a potential importance of fast-endophilin-mediated endocytosis (FEME^21^) in this tumor type.

**Figure 4.**
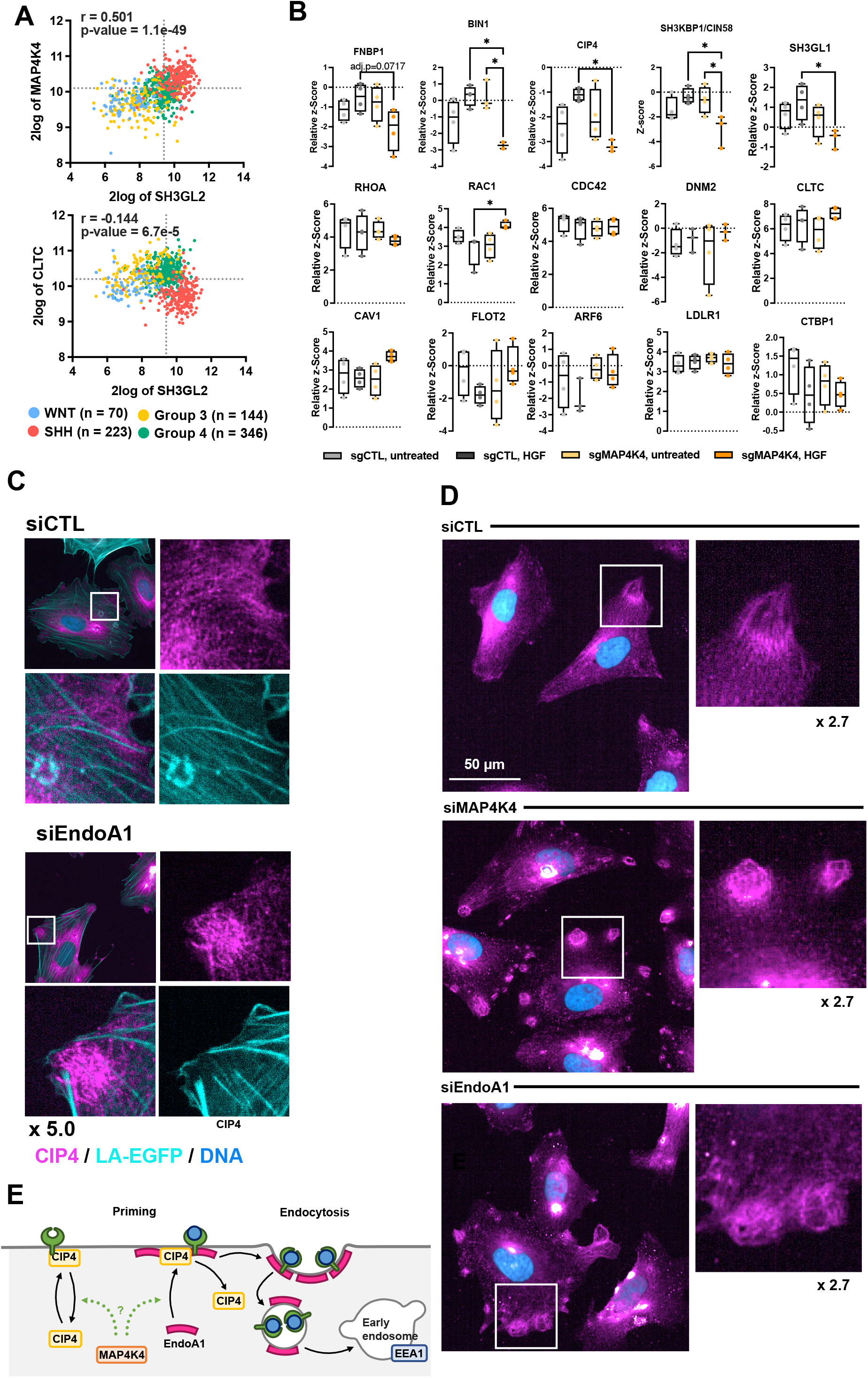
MAP4K4 controls CIP4 PM association and subcellular localization. **(A**) Two gene analyses of SH3GL2, MAP4K4 and CLTC expression across the four MB subgroups. Correlation coefficient and adjusted p-values are shown. **(B)** Quantification of PM association of regulators of FEME priming and endocytosis effectors. Box plots display normalized z-score values extracted from all-group comparison of MS analysis. N = 4 independent experiments, min to max, * = p<0.05 of one-way ANOVA. **(C)** Confocal microscopy analysis of endogenous CIP4 localization in siCTL or siEndoA1 transfected ONS-76 cells. Cyan: LA-EGFP (F-actin), magenta: CIP4, blue: Hoechst (DNA). **(D)** Confocal microscopy analysis of CIP4 subcellular localization in siCTL, siMAP4K4 and siEndoA1 transfected cells. Magenta: CIP4, blue: Hoechst. **(E)** Model for MAP4K4 regulation of the fast endophilin mediated endocytosis. Under control conditions, during priming: CIP4 is recruited to the plasma membrane (PM). PM-associated CIP4 recruits EndoA1 to the plasma membrane. After receptor activation, endophilins initiate FEME and promote the internalization of the carrier and trafficking towards early endosomes.

### MAP4K4 controls PM association and subcellular distribution of FEME regulators

To explore whether MAP4K4 could be involved in PM localization of FEME components such as CIN85/SH3KBP1, CIP4, BIN1 and FBP17, four regulators of FEME priming^55^, we compared their abundance in the different surface proteome samples we generated (Tables 1 & 2). We found that HGF stimulation caused a small increase in PM association of these proteins and of EndoA2 in control cells (Fig. 4B). In contrast, PM association of CIN85/SH3KBP1, CIP4, BIN1 and EndoA2 decreased significantly in HGF-stimulated sgMAP4K4 cells, indicating that MAP4K4 controls membrane association of these proteins in cells with activated c-MET. We could not detect EndoA1 and EndoA3 in pour proteomic samples. CIN85/SH3KBP1, CIP4 and FNBP1/FBP17 are considered critical mediators of FEME by priming the PM for endophilin A clustering^55^. PM enrichment of these proteins in a MAP4K4-dependent manner in HGF-stimulated cells could thus indicate a link between MAP4K4 and FEME priming. In cells, CIP4 is detectable associated with dotted and filamentous structures that do not co-localize with F-actin (Fig. 4C, D). The depletion of MAP4K4 alters the subcellular localization of CIP4 and causes CIP4 to accumulate in circular patches near cellular protrusions (Fig. 4D). Depletion of EndoA1 phenocopies CIP4 accumulation in these patches (Fig. 4D). Interestingly, we also observed the early endosome marker EEA1 in similar patches in control but not in MAP4K4-or EndoA1-depleted cells (Fig. S5). Whereas this potential colocalization remains to be further investigated, we observed that depletion of EndoA1 or MAP4K4 led to a 20% reduction of EEA1 puncta per cell in a HGF stimulation-independent manner. This indicates that both EndoA1 and MAP4K4 enable formation of early endosomes and further corroborate the shared functionality of the two proteins in endocytosis regulation. Together, these results indicate the implication of MAP4K4 in membrane recruitment, cortical localization and organization of key regulators of FEME in HGF stimulated cells, thereby possibly facilitating an effective FEME sequence^55^ (Fig. 4E).

### EndoA family proteins are required for cell motility and tissue invasion

To determine whether endocytosis pathways (Fig. S6A) are involved in HGF-induced migration control, we assessed the contributions of endocytic pathway components to the HGF-induced invasive phenotype. We used 25 siRNAs targeting 16 essential components of 8 different endocytic pathways (Fig. S6B). Downregulation efficacy of the siRNAs was confirmed by RT-qPCR (Fig. S6C) and HGF induced transwell migration assessed using Boyden chamber assays (Fig. 5A). Consistent with previous studies^13, 14^, HGF stimulation increased the number of transmigrated cells (∼ 2.7-fold increase). Depletion of dynamin 1 and 2 (DNM1 and DNM2) abrogated HGF-induced migration (Fig. 5A), confirming a general role of endocytosis in migration control. Interestingly, depletion of Endophilin A proteins completely abrogated (SH3GL2/EndoA1) or drastically reduced (SH3GL1/EndoA2 and SH3GL3/EndoA3) HGF-induced cell migration (Fig. 5A). In contrast, depletion of clathrin or caveolin, two essential components of endocytic pathways for GF signaling^56^, did not prevent or reduce HGF-induced DAOY cell migration. Similarly, depletion of two regulators of micropinocytosis, CTBP1 and RAC1^57^, did also not block HGF-induced migration. Both non-transfected and siCTL transfected cells showed similar levels of proliferation up to 72 hours, while depletion of SH3GL1/2/3 and DNM1 exhibited even moderately increased levels of proliferation (Fig. S6D). These data exclude a proliferation defect as the underlying cause of reduced transwell migration in siSH3GL1/2/3- or siDNM1-transfected cells and indicate the implication of EndoA1 and DNM1 in cell migration control.

**Figure 5.**
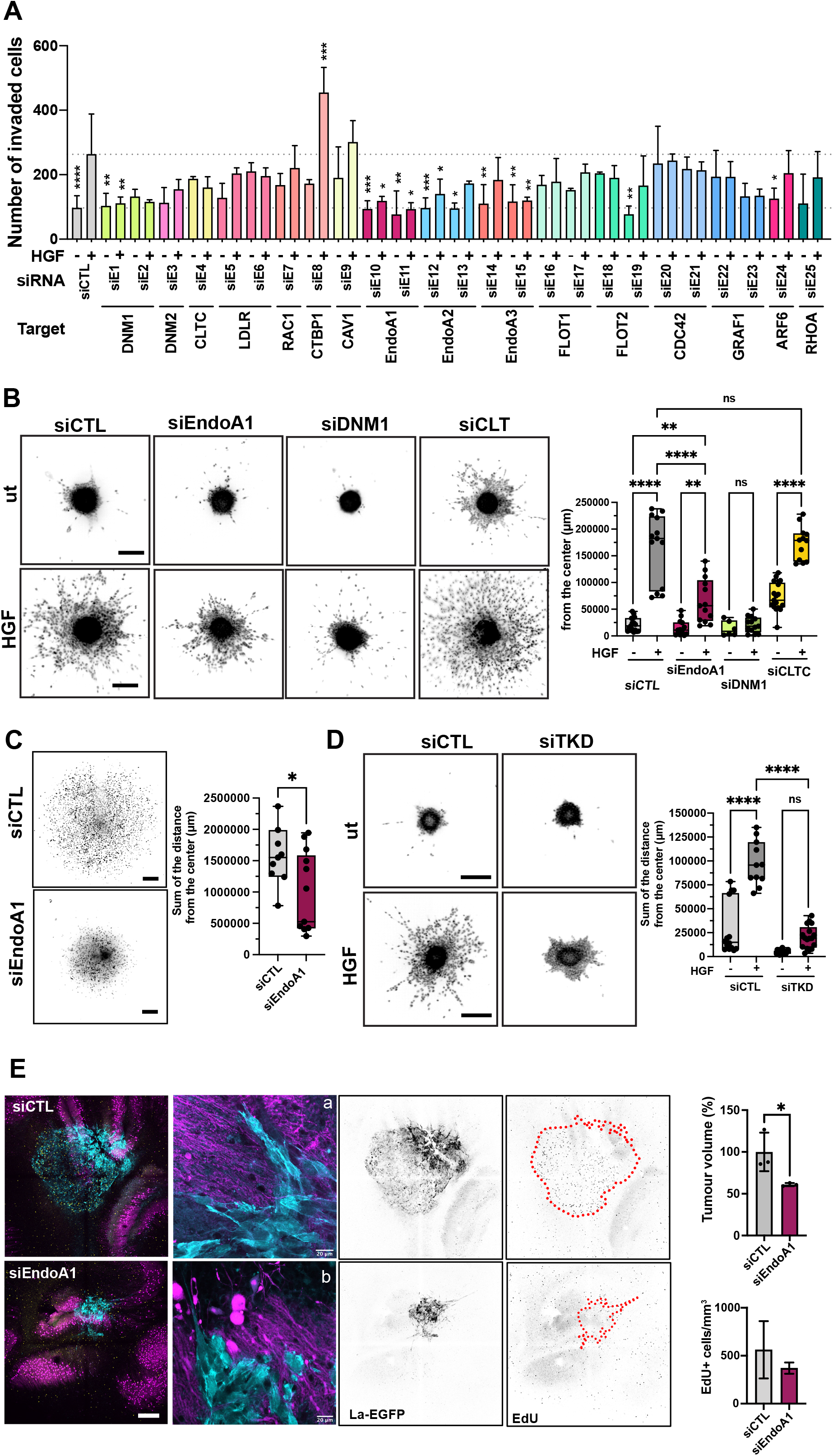
EndoA1 is required for migration and tissue invasion. **(A)** Boyden chamber transwell migration assay using siRNA transfected DAOY cells. siRNAs as indicated. Significance is calculated from the comparison of each individual sample with the HGF-stimulated siCTL sample. N = 3-4 independent experiments, mean & SD, * = p<0.05, ** = p<0.01, *** = p<0.001 of one-way ANOVA. **(B)** SIA analysis of EndoA1 implication in collagen I invasion. Left: Inverted greyscale images of Hoechst-stained nuclei of DAOY spheroids ± HGF (20 ng/ml) at endpoint. siRNA transfections as indicated. Scale bar = 300 µm. Right: Quantification of the total distance of invasion as sum of all distances measured from the center of spheroids as shown in A. Each dot represents 1 spheroid, n = 3 independent experiments, mean & SD, ** = p<0.01, **** = p<0.0001 of one-way ANOVA. **(C)** Left: Inverted greyscale images of Hoechst-stained nuclei of ONS-76 spheroids at end point of SIA. Scale bar = 300 µm. Right: Quantification of the invasion as sum of the distances as in B, * = p<0.05 of unpaired t-test. **(D)** Left: Inverted greyscale images of Hoechst-stained nuclei of DAOY cells transfected with three siRNAs targeting EndoA1, A2 and A3 (TKD) ± HGF (20 ng/ml) at end point of SIA. Scale bar = 300 µm. Right: Quantification of the invasion as sum of the distances as in B, **** = p<0.0001 of one-way ANOVA. Scale bar in all representative images = 300 µm. **(E)** Left: Maximum intensity projection of representative confocal sections of OCSC implanted with transfected ONS-76 spheroids 48h after implantation. Cyan: Lifeact-EGFP; Magenta: calbindin (Purkinje cells). Right: Quantification of tumor volume and proportion of EdU-positive cells/area. N = 3, technical replicates, mean & SD, * = p<0.05, unpaired t-test. Scale bar = 400 µm. a and b are higher magnification images of ONS-76 tumor cells at the invasion front. Cyan: Actin cytoskeleton of tumor cells, magenta: Purkinje cells.

### Endophilin A1 is required for 3D collagen I matrix invasion

We next tested whether depletion of EndoA1, DNM1, or clathrin heavy chain (CLTC) also affects 3D collagen I invasiveness of MB cells (Fig. 5B, S6E, F). HGF caused a robust increase in the number of cells disseminating into the collagen I matrix and increased the distance of invasion (∼ 6.8-fold increase compared to untreated control). Depletion of EndoA1 or DNM1 significantly reduced HGF-induced collagen I invasion, whereas depletion of CLTC had no effect, indicating that EndoA1 functions in clathrin-independent endocytosis. Consistent with a conserved role of EndoA1 in migration control, depletion of EndoA1 also reduced (∼ 1.6-fold decrease) collagen I invasion of ONS-76 cells (Fig. 5C). Depletion of MAP4K4 caused also some, albeit insignificant, reduction in EndoA1 expression (Fig. S6F). To determine whether a compensatory mechanism mediated by EndoA2 or EndoA3^24^ may be active in our cell model, we depleted all Endophilin-A proteins simultaneously (TKD). We found that endophilin TKD abrogates HGF-induced cell migration in DAOY cells moderately more effectively than siEndoA1 alone (Fig. 5D). We concluded that EndoA2 or EndoA3 contribute to migration control in MB cells, and that EndoA1 is the most relevant Endophilin in our cell model. In conclusion, we found that EndoA1 mediates a migratory phenotype in MB cells, possibly through a mechanism involving clathrin-independent endocytosis.

### Endophilin-A1 depletion reduces tissue invasion

We next assessed the role of EndoA1 in tumor growth and invasion control in the tissue context with tumor-cerebellar slice co-culture experiments using ONS-76-LA-EGFP cells. Depletion of EndoA1 markedly reduced cerebellum tissue infiltration and prevented the formation of streams of invading cells, which are highly characteristic for this type of MB tumor cell (Fig. 5E, panel a). Individual EndoA1-depleted tumor cells still displayed invasion at the border of the spheres. However, the morphology of these cells appeared more rounded, less mesenchymal compared to control cells (Fig. 5, panel b). Although morphology of the collectivity of the invading cells varied between samples, the considerably reduced invasiveness of EndoA1-depleted cells was apparent (Fig. 5E, S6G), while proliferation was not significantly affected *in vitro* (Fig. S6D) and *ex vivo* (Fig. 5E). Taken together, these data indicate that EndoA1 contributes to the invasive behavior of the tumor cells in the cerebellar tissue without affecting proliferation.

## Discussion

We found that HGF-induced c-MET activation alters the abundance of transmembrane and plasma membrane-associated proteins in MB cells. The abundance of many of these proteins is reduced by HGF-c-MET activation, indicating either HGF-dependent internalization or attenuated recycling of PM-associated proteins. HGF-induced changes in PM association are increased in MAP4K4-depleted cells and significantly different to control cells for a number of these proteins. Proteins belonging to this group of MAP4K4-regulated proteins are the adhesion receptor CD155 and regulators of fast-endophilin-mediated endocytosis. We show that CD155 can contribute to the oncogenic functions of the MB tumor cells by enabling migration, proliferation and tumor cell growth in the brain tissue. We furthermore identified the fast-endophilin-mediated endocytosis effector protein EndoA1 as a mediator for HGF-induced collagen invasion *in vitro* and for tissue invasion *ex vivo*. The implication of MAP4K4 in PM association of regulators, effectors and targets of endocytic turnover provides a novel conceptual basis for strategies to target the oncogenic phenotype of MB and other RTK-driven solid tumors.

PM-associated proteins were enriched by biotin labeling prior to mass-spectrometry analysis. This methodology is expected to label predominately extracellularly bound or anchored proteins, transmembrane proteins, and proteins associated with endocytic vesicles. However, internalized biotin may also lead to the purification of intracellular proteins not normally associated with the plasma membrane. To exclude non-PM proteins, we either used the TMHMM algorithm to predict transmembrane domains in proteins^29^ or a data-based filtering approach, yielding two datasets. Former detects surface proteins. However, it also predicts transmembrane domains of proteins associated with other membrane compartments such as the Golgi or ER membranes. The data-based filtering approach selects proteins based on published subcellular localization^30^. This approach is not specifically enriching for proteins with transmembrane domains, and it may include non-PM proteins complexed with PM-associated proteins. The TMHMM and data-based annotation approach selected 21% and 8% of all proteins detected in the initial MS analyses, respectively. The TMHMM approach was highly efficient for determining MAP4K4 impact on PM-association of transporters and channels, which all contain a TM domain. The data-based annotation approach allows for the comparative analysis of the variations in PM association in all the conditions tested using pathway enrichment analysis.

MAP4K4 implication in HGF-induced internalization of membrane transport proteins and drug transporters^31^ indicated a potential therapeutic benefit of targeting this function of MAP4K4 during chemotherapy, which remains one of the first-line treatments for cancer patients. We hypothesized that indirect modulation of drug uptake or extrusion mechanisms could improve drug efficacy. MAP4K4 depletion moderately decreased the sensitivity of DAOY cells to Lomustine, but it had no effect on etoposide treatment efficacy. Although of potential interest, our data are inconclusive and susceptibilities to additional chemotherapeutic drugs such as cisplatin, which is internalized through LRRC8A^58^, should be tested in MAP4K4 depleted cells. Consistent with a potential implication of MAP4K4 in sensitization of tumor cells to chemotherapy, MAP4K4 was recently found to reduce the sensitivity of cervical cancer cells to cisplatin treatment via an autophagy-controlled mechanism^59^. The disturbed cell-surface localization of ion channels and transporters could also modify the overall ion homeostasis of the cell. This notion is supported by the subtle alterations in electrical currents we observed in MAP4K4-depleted cells, and it is consistent with previous studies that demonstrated altered electrical conductance properties in tumors compared to healthy tissues^37, 60, 61^. We found several ion channels susceptible to MAP4K4 regulation and observed an increased current in MAP4K4-depleted cells, together indicating indirect MAP4K4 control of ion homeostasis via regulated ion channel distribution. The lowered potential in MAP4K4-depleted cells after HGF stimulation could be the consequence of a reduced influx of cations, increased efflux of cations, or increased influx of anions. We observed increased PM-association of potassium channels and potassium transport-associated molecules including the P-type cation transporter ATP12A, the Ca^2+^-dependent channels ATP1A3 and KCNMA1 in MAP4K4-depleted cells after HGF stimulation. Thus, MAP4K4 could maintain optimal polarization in HGF-stimulated cells by controlling potassium efflux rates through regulating PM-association of K^+^ transporters. However, whether differential PM association of potassium channels and de-regulated potassium fluxes are at the origin of the electrical disbalance observed in MAP4K4-depleted cells and contribute to the MAP4K4-associated pro-migratory phenotype remains to be determined.

PM-association of CD155 is reduced in MAP4K4-depleted cells that are stimulated with HGF. In glioma cells, CD155 mediates adhesion to the extracellular matrix protein vitronectin, promotes FA turnover and FA signaling towards SRC, paxillin and p130CAS, and thereby contributes to tissue invasion^62^. MAP4K4 contributes to FAK activation at the leading edge of migrating cells^14^, and our data herein indicate a possibly indirect regulation of this process via the control of CD155 surface expression. Our observation of a more rounded, less mesenchymal morphology in CD155-depleted cells has been described previously in glioma cells^14^, and studies in triple-negative aggressive breast cancer cells demonstrated CD155 contribution to the mesenchymal cell state^63^. Depletion of CD155 in these cells triggered mesenchymal to epithelial transition and repressed migration and invasion *in vitro* and *in vivo*., suggesting that CD155 expression in MB cells could also contribute to migration control indirectly via transcriptional reprogramming. CD155 also represents an attractive therapeutic target due to its ability to inhibit NK cell and CD8^+^ T-cell activity, thereby contributing to immune evasion of tumor cells^64, 65^. Kinase-controlled modulation of CD155 surface expression -as we found herein – could enable this process as rather the abundance, and not the absolute absence or presence, of CD155 at the surface of the cell is critical for its immunomodulatory capabilities^49^. Under physiological conditions, CD155 is expressed at low levels, where a balance between its immune activating and inhibitory functions maintains the normal function of immune cells. Thus, the MAP4K4-dependent maintenance of surface expression of CD155 on growth factor-activated cells could not only affect the migratory potential of the cells but also locally (in the tumor microenvironment) repress immune system activation.

Our findings point towards a role of fast-endophilin-mediated endocytosis (FEME) through EndoA1 in the control of the invasive phenotype of MB cells. EndoA1 and A3 are highly expressed in SHH MB compared to other brain tumors and their expression correlates positively with MAP4K4 in this tumor type. This contrasts with clathrin, which correlates negatively with MAP4K4 in SHH MB and with the lack of clear evidence that clathrin-mediated endocytosis is necessary for migration control in our cell models. The role of endophilins in cancer progression is complex and controversial, as endophilins can both promote and reduce cancer cell migration, depending on tumor type^24, 55, 66^. We previously found that 5-(*N*-ethyl-*N*-isopropyl)amiloride (EIPA), a micropinocytosis inhibitor^67^, blocks HGF-induced invasion^14^. However, depletion of RAC1 or CTBP1, two upstream activators of micropinocytosis, did not significantly impact HGF-induced motility. This suggests that the pronounced effect of EIPA could be due to the endocytosis-unrelated effects of this drug on ion transport, intracellular pH and cytoskeleton regulation, thus affecting FEME indirectly^24, 55^. FEME is involved in the endocytosis of activated receptors at the leading edge of migrating cells^68^ and fast and ultrafast endocytosis are important for chemotaxis^69^. FEME could be involved in the internalization of c-MET, thereby controlling signal transmission and persistence of this receptor^24^. Our study identified a possible link between MAP4K4 function and FEME through the regulated PM-association of the FEME priming factors SH3KBP1, CIP4 and FBP17. The observation that MAP4K4 regulates CIP4 PM-association and its accumulation in cortical patches provides circumstantial evidence of spatial regulation of CIP4 by MAP4K4. The early endosome marker EEA1 also localizes to these patches, and it is thus possible that one aspect of FEME processing towards early endosomes occurs preferentially there. Depletion of MAP4K4 prevented EEA1 recruitment to these patches, led to an overall decrease in EEA1-positive vesicles and caused accumulation of CIP4 in this location instead. These phenomena are quite exactly phenocopied by EndoA1 depletion. It is thus possible that MAP4K4 controls FEME by orchestrating a step between FEME priming and trafficking towards early endosomes. Our working model is that MAP4K4 orchestrates PM association of CIP4 from a non-PM-associated subcellular compartment to the “CIP4 patch” in close proximity of or associated with the PM, where also early endosomes enrich. Upon HGF stimulation, CIP4 translocates to the PM, where it is rapidly turned over through EndoA1-dependent FEME before it is recycled back to the PM. Without MAP4K4 function, CIP4 remains associated with the patches, hence the overall amount of PM associated CIP4 increases. Some key questions remain: (i) Does the MAP4K4-dependent mechanism of action involve phosphorylation of one of the components involved in FEME, (ii) is MAP4K4 regulation of CIP4 involved in CIP4-dependent recruitment of Endophilin-A1 to the plasma membrane and (iii) does MAP4K4 control of FEME priming involve the activation of CDC42 downstream of GEFs and GAPs. Elucidation of these mechanisms will be essential to explore therapeutic targeting strategies for specifically repressing MAP4K4 controlled pro-migratory endocytosis.

In conclusion, we found that the HGF-c-MET-MAP4K4 axis regulates PM association and surface expression of proteins affecting different pro-oncogenic mechanisms, including chemotherapy sensitivity, ion homeostasis and cell migration and proliferation. Furthermore, our study indicates the contribution of endophilin A proteins in invasion control downstream of MAP4K4. It thus highlights kinase regulation of the PM proteomic composition as a novel mechanism involved in establishing and maintaining the oncogenic phenotype in MB and other solid tumors. The study provides a conceptual basis for further analysis of the dynamic adaptation of tumor cells to environmental cues by plasma membrane proteome modification. Finally, targeting MAP4K4 controlled endocytic activity could represent a novel druggable vulnerability in medulloblastoma cells to restrict tumor growth and dissemination.

## Materials and Methods

### Cell lines and culture conditions

DAOY human MB cells were purchased from the American Type Culture Collection (ATCC, Rockville, MD, USA). ONS-76 cells were generously provided by Michael Taylor (SickKids, Canada). DAOY and DAOY-LA-EGFP cells were cultured in iMEM complemented with 10% FBS. Medium without FBS is used for starvation and further referred to as serum-free medium (SFM). ONS-76-LA-EGFP cells were cultured in RPMI 1640 complemented with 10% FBS. Medium without FBS is used for starvation. DAOY and ONS-76 Lifeact-enhanced green fluorescent protein (LA-EGFP) cells were produced by lentiviral transduction of DAOY cells with pLenti-LA-EGFP. Cell line authentication and cross-contamination testing were performed by Multiplexion GmbH (Heidelberg, Germany) by single nucleotide polymorphism (SNP) profiling.

### Animals

Pregnant Wild type C57BL/6JRj females were purchased from Janvier Labs and were kept in the animal facilities of the University of Zürich Laboratory Animal Centre. Mouse protocols for organotypic brain slice culture were approved by the Veterinary Office of the Canton Zürich.

### Generation of CRISPR/Cas9-mediated MAP4K4 knockout cells

MAP4K4 gene-specific single-guide RNA (sgRNA) were designed using Synthego CRISPR design online tool (https://design.synthego.com). Oligos were synthesized by Microsynth (Balgach, Switzerland) and cloned into LentiCRISPRv2 transfer plasmid (Addgene, 52961) with a single tube restriction and ligation method as described by McComb *et al*.^70^. Production of lentiviral vectors and cell transduction was performed as described previously^14^. The efficiency of the knockouts was tested by immunoblot. Only the guide with the highest efficiency has been selected for further experiments (sgMAP4K4#1, exon 7, GGGCGGAGAAATACGTTCAT).

### Cell-surface labelling

DAOY cells were prepared in four T75 flasks to reach a 70% confluency (per condition and replicates). Standard medium was then replaced with SFM. After 24 hours, cells were 85% confluent, and media were replaced with either pre-warmed SFM or pre-warmed SFM supplemented with 20 ng/mL HGF (PeproTech, 100-39) for 30 min. EZ-Link-Sulfo-NHS-SS-biotin kit (Thermo Scientific, 89881) was used to surface label DAOY_sgCTL and DAOY_sgMAP4K4 cells. In brief, 10 ml EZ-Link-Sulfo-NHS-SS (0.25 mg/ml, diluted in ice-cold PBS) was added at 4°C for 30 minutes. Biotinylation reaction was terminated by addition 500 µl of provided quenching solution. Cells from four T75 were collected in a unique tube and washed three times with 10 ml ice-cold TBS (700g, 3 minutes centrifugations). Cell pellets were lysed on ice for 30 minutes with one cycle of sonication (Sonoplus, HD2070) and 5 seconds vortexing every 5 minutes. Insoluble debris pelleted using centrifugation at 10’000 g for 2 minutes. Biotinylated proteins were affinity-purified from lysates using columns containing NeutrAvidin Agarose slurry for one hour at room temperature with end-over-end mixing. After 3 washing steps with lysis buffer, bound proteins were eluted using sample buffer (BioRad, 1610747), complemented with 50 mM DTT, for one hour at room temperature with end-over-end mixing. Purified lysates were kept at -20°C for up to 1 day before Mass spectrometry analysis.

Samples were prepared for mass-spectrometry (MS) by using the iST Kit (PreOmics, Germany) according to a modified protocol. In brief, proteins were precipitated with a 10% TCA, solubilized in 50 µl iST Kit lysis buffer, boiled at 95°C for 10 minutes and processed with High Intensity Focused Ultrasound (HIFU) for 30 s, with ultrasonic amplitude set to 85%. Proteins were quantified using Qubit, and 50 µg of the samples were for digestion using 50 µl of the iST Kit digestion solution. After 60 min f incubation at 37°C, the digestion was stopped with 100 µl of iST Kit Stop solution. The non-bound components of the solutions in the cartridge were removed by centrifugation at 3800 x g, while the peptides were retained by the iST-filter. Finally, the peptides were washed, eluted, dried and re-solubilized in 20 µl iST Kit LC-Load buffer for MS-Analysis.

### Liquid chromatography-mass spectrometry analysis

MS analysis was performed on a Q Exactive HF-X mass spectrometer (Thermo Scientific) equipped with a Digital PicoView source (New Objective) and coupled to a M-Class UPLC (Waters). Solvent composition at the two channels was 0.1% formic acid for channel A and 0.1% formic acid, 99.9% acetonitrile for channel B. For each sample, 1 μl of peptides were loaded on a commercial MZ Symmetry C18 Trap Column (100Å, 5 µm, 180 µm x 20 mm, Waters) followed by nanoEase MZ C18 HSS T3 Column (100Å, 1.8 µm, 75 µm x 250 mm, Waters). The peptides were eluted at a flow rate of 300 nl/min by a gradient from 8 to 27% B in 85 min, 35% B in 5 min and 80% B in 1 min. Samples were acquired in a randomized order. The mass spectrometer was operated in data-dependent mode (DDA), acquiring a full-scan MS spectrum (350−1’400 m/z) at a resolution of 120’000 at 200 m/z after accumulation to a target value of 3’000’000, followed by HCD (higher-energy collision dissociation) fragmentation on the twenty most intense signals per cycle. HCD spectra were acquired at a resolution of 15’000 using a normalized collision energy of 25 and a maximum injection time of 22 ms. The automatic gain control (AGC) was set to 100’000 ions. Charge state screening was enabled. Singly, unassigned, and charge states higher than seven were rejected. Only precursors with intensity above 110’000 were selected for MS/MS. Precursor masses previously selected for MS/MS measurement were excluded from further selection for 30 s, and the exclusion window was set at 10 ppm. The samples were acquired using internal lock mass calibration on m/z 371.1012 and 445.1200. The mass spectrometry proteomics data were handled using the local laboratory information management system (LIMS)^71^. The mass spectrometry proteomics data have been deposited to the ProteomeXchange Consortium via the PRIDE^72^ partner repository with the dataset identifier PXD030597.

### Protein identification and label-free protein quantification

The acquired raw MS data were processed by MaxQuant (version 1.6.2.3), followed by protein identification using the integrated Andromeda search engine^73^. Spectra were searched against a Swissprot Homo sapiens reference proteome (taxonomy 9606, version from 2016-12-09), concatenated to its reversed decoyed fasta database and common protein contaminants. Carbamidomethylation of cysteine was set as fixed modification, while methionine oxidation and N-terminal protein acetylation were set as variables. Enzyme specificity was set to trypsin/P, allowing a minimal peptide length of 7 amino acids and a maximum of two missed cleavages. MaxQuant Orbitrap default search settings were used. The maximum false discovery rate (FDR) was set to 0.01 for peptides and 0.05 for proteins. Label-free quantification was enabled and a 2-minute window for the match between runs was applied. In the MaxQuant experimental design template, each file is kept separate in the experimental design to obtain individual quantitative values. Protein fold changes were computed based on Intensity values reported in the proteinGroups.txt file. A set of functions implemented in the R package SRMService (http://github.com/protViz/SRMService) was used to filter for proteins with two or more peptides allowing for a maximum of four missing values, and to normalize the data with a modified robust z-score transformation and to compute p-values using the t-test with pooled variance. If all protein measurements are missing in one of the conditions, a pseudo fold change was computed replacing the missing group average by the mean of 10% smallest protein intensities in that condition. For data visualization and normalization across all samples, all group comparison correspond has been processed by statistical analysis of all sgMAP4K4 samples (starved and stimulated) versus all sgCTL samples (starved and stimulated).

### Chemotherapy sensitivity analysis

400 DAOY sgCTL or sgMAP4K4 were seeded per well in 384 well microplate and incubated 24 hours at 37°C. Media were either replaced with low-serum media (1% FBS) or fresh standard culture media (10% FBS) and incubated for another 24 hours at 37°C. Cells were treated with either low-serum media or low-serum media supplemented with 20 ng/mL HGF (PeproTech, 100-39) or by standard culture media supplemented with Lomustine (Selleckchem, S1840, 5 µM to 100 µM) or Etoposide (Selleckchem, S1225, 1 nM to 10 µM). Cell proliferation was quantified after 48 h of incubation using cell proliferation reagent WST-1 (Roche, 11644807001) following the manufacturer’s instructions. Sample absorbance was measured against the background using a microplate reader (Biotek Instruments, Cytation 3) and statistically analyzed using Prism software (GraphPad).

### Primary tumor gene expression analysis

Gene expression data were obtained from the R2 genomics and visualization platform (https://r2.amc.nl). Berchtold – 172 – MAS5.0 – u133p2; Normal cerebellum – Roth – 9 – MAS5.0 – u133p2; Tumor Medulloblastoma (SHH) – Pfister – 73 – MAS5.0 – u133p2; Tumor Medulloblastoma – Hsieh – 31 – MAS5.0 – u133p2; Tumor Medulloblastoma – Pfister – 273 – MAS5.0 – u133p2; Tumor Medulloblastoma public – Delattre – 57 – MAS5.0 – u133p2; Tumor Medulloblastoma PloS One – Kool – 62 – MAS5.0 – u133p2; Tumor Medulloblastoma – Gilbertson – 76 – MAS5.0 – u133p2 were used to analyze CLIC1 expression in Medulloblastoma versus normal brain tissues. The Tumor Medulloblastoma – Cavalli – 763 – rna_sketch – hugene11t dataset with 763 primary medulloblastoma samples was used for Kaplan Meier survival analysis relative to CLIC1 mRNA expression levels. The Tumor Medulloblastoma – Cavalli – 763 – rna_sketch – hugene11t dataset with 763 primary medulloblastoma samples was used. Molecular pathways correlating with MAP4K4 gene expression have been determine using the R2 genomics and visualization platform built-in KEGG PathwayFinder by Gene correlation tool. Transcriptomic and proteomic data were obtained from the ProTrack pediatric brain tumor database of the Clinical proteomic Tumor Analysis Consortium (CPTAC) and Children’s Brain Tumor Tissue Consortium (CBTTC).

### Patch-clamp analysis

DAOY sgCTL or sgMAP4K4 were seeded in a 35 mm^2^ dish to reach 20% confluency. After 6 hours of incubation in standard culture condition, the medium was replaced with starvation medium. The next day, cells were treated with either SFM or SFM supplemented with 20 ng/ml HGF for 30 min. Cells were kept in the patch chamber for 30 min. Whole-cell recordings at room temperature were performed on DAOY sgCTL and sgMAP4K4 cells using patch pipettes of ∼7 MΩ. The patch pipette solution contained (in mM) 95 K-gluconate, 30 KCl, 4.8 Na_2_HPO_4_, 1.2 NaH_2_PO_4_, 5 glucose, 2.38 MgCl_2_, 0.76 CaCl_2_, 1 EGTA, 3 K-ATP. The bath solution contained (in mM) 142 NaCl, 1.8 MgCl_2_, 1.8 CaCl_2_, 10 HEPES, 3 KCl. Compensations and measurements were done within 30 seconds after obtaining whole-cell to avoid drastic changes in cytoplasmic ionic concentration.

### Gene annotation and pathway enrichment analysis

Gene annotation and pathway enrichment analysis on the surface-proteomics results were performed using the Metascape webtool (https://metascape.org) developed by Zhou *et al.*^39^. Transmembrane proteins were annotated using the TMHMM prediction algorithm^29^. We performed pathway enrichment using all pathway enrichment categories, standard settings and using all proteins identified by mass-spectrometry as background genes.

### siRNA transfections

100’000 DAOY or ONS-76 cells were seeded per well in 6-well plates. The following day, cells were transfected with 10 nM siRNAs using Lipofectamine RNAiMAX Transfection Reagent (Invitrogen, 13778075) plus. siRNA used are referenced in Table S1. After 6 hours, the media were changed, and cells were kept in culture overnight for downstream analyses.

### RT-qPCR analysis of gene expression

100’000 siRNA transfected DAOY cells (24 h after transfection) were seeded per well in 6 well plates and incubated overnight. For quantitative real-time PCR (RT-qPCR) analysis of target genes, total RNA was isolated using Qiagen Rneasy Mini Kit (Qiagen, 74106). One µg of mRNA was converted to cDNA using the high-capacity cDNA Reverse Transcription Kit (Applied Biosystems, 4368813). RT-qPCR was performed using PowerUp Syber Green (Applied Biosystems, A25776) under conditions optimized for the ABI7900HT instrument. The primers used are referenced in table S2. The ΔΔCT method was used to calculate the relative gene expression of each gene of interest.

### Flow cytometry analysis of CD155 surface expression

300’000 siRNA transfected DAOY cells were seeded 24h after transfection per well in low adhesion 6 well plate (Greiner Bio-one, 657970) and incubated for 6 h in SFM. Cells were then treated with either pre-warmed SFM or SFM supplemented with 20 ng/ml HGF for 30 min. After stimulation, cells were fixed by adding paraformaldehyde pre-warmed to 37°C to a final concentration of 4% and incubated for 20 min at 37°C. After one wash with PBS supplemented with 2% FBS, cells were stained with PE-labelled anti-human CD155 (BioLegend, 337610, 1:300), corresponding isotype control antibody (BioLegend, 400114, 1:300), or left unstained for 20 min on ice. Samples were washed with PBS supplemented with 2% FBS, resuspended in PBS and acquired using BD LSRFortessa flow cytometer (BD Bioscience). Fluorescence levels were analyzed using FlowJo software (BD Bioscience).

### Spheroid invasion assay

2’500 siRNA transfected cells were seeded 24h after transfection per well in 100 µl of standard growth media in cell-repellent 96 well microplates (Greiner Bio-one, 650790). Plates were incubated 24 hours at 37°C in 5% CO_2_ to allow spheroid formation. 70 µl of culture media were removed from each well and the remaining medium containing the spheroids was complemented with a collagen solution (2.5 mg/ml Pure Col Collagen I (Advanced Biomatrix), DMEM 1x (Sigma, D2429) and 0.4% Sodium bicarbonate (Sigma, S8761)). After collagen I polymerization (∼ 2 hours), fresh 100 µl SFM +/- HGF 20 ng/ml for DAOY spheroids or standard culture media for ONS-76 spheroids was added per well. Cells were incubated at 37°C for 16 hours until fixation with 4% paraformaldehyde. A 1:5000 dilution of Hoechst (Sigma Aldrich, B2883) was added to stain DNA for nuclei visualization. Images of spheroids and invaded cells were acquired on an Operetta CLS High-Content Analysis System (PerkinElmer, HH16000000) at 5x magnification using the Dapi (405 nm) channel. Spheroids were localized in the well by acquiring 3 z-planes with 250 µm z-spacing. Localized spheroids were re-scanned by acquiring 16Hoechst-stained z-planes with 25 µm z-spacing and a maximum intensity projection of each spheroid was computed and used for further analysis. Using Harmony software (PerkinElmer), spheroids and invading cells were delineated based on fluorescence threshold. The distance from the center of the spheroid was calculated, and the sum of the cell invasion was determined as the sum of all individual values acquired per well.

### 3D-Cell viability/proliferation (Cell TiterGlo) assay

Cell viability was determined using CellTiter-Glo® 3D cell viability assay (#G9242, #G9682, Promega). 500 siRNA transfected cells/25µL were seeded in U-low adhesion (#4516, Corning) 384-well plate. After 48h, the CellTiter-Glo® 3D reagent was added (volume/volume) following the manufacturer’s instructions. Plates were incubated at room temperature under agitation for 30 min and luminescence representing the number of viable cells was quantified with a Cytation 3 imaging reader (BioTek®).

### Cell proliferation assay

1’000 DAOY cells were seeded in 96 well plates (Greiner Bio-One, 655087). The following day, cells were transfected according to the protocol described above. Six hours after transfection, medium was replaced with serum-free medium. For timepoint 0, WST reagent was added to the corresponding well and incubated 30 min at 37°C. Absorbance at 440 nm was measured using a microplate reader (Biotek Instruments, Cytation 3). 20 ng/ml HGF was added to remaining timepoints and incubated for 24, 48, and 72 hours. At the end of each timepoint, WST reagent was added to the corresponding well and absorbance measured. Each day media was replaced with fresh serum-free medium supplemented with HGF. Changes in absorbance over time were analyzed using Prism 9 software (GraphPad).

### Single-cell motility assay

1’000 siRNA transfected ONS-76-LA-EGFP cells were seeded in 96 well microplates (Greiner Bio-one, 655098) previously coated with Pure Col Collagen I (Advanced BioMatrix, 5005) at 10 µg.cm^-2^. After 24 h incubation under standard culture conditions, cell motility was acquired using temperature (37°C) and CO_2_ (5%) controlled Operetta CLS High-Content Analysis System (PerkinElmer, HH16000000) (non-confocal, 20x objective, 4 fields per well, 3 min intervals, 3 h acquisition). Cell segmentation was performed using the LA-EGFP channel. Cell speed, area, circularity, and the width/length ratio were calculated over time using Harmony software (PerkinElmer).

### *Ex vivo* Organotypic Cerebellum Slice Culture (OCSC)

*Ex vivo* Organotypic Cerebellum Slice Culture was carried out essentially as described previously^56^. Wild type C57BL/6JRj mice were sacrificed at postnatal day 8-10. Cerebella were dissected and kept in ice-cold Geys balanced salt solution containing kynurenic acid (GBSSK) and then embedded in 2% low melting point agarose gel. 350 µm thick sections were cut using a vibratome (Leica, VT1200S) and transferred on inserts (Merck Millipore, PICM03050) for further *in vitro* culture. Slices were kept in culture and monitored for 15 days, and media were changed daily for the first week and once in two days thereafter. Spheroids of DAOY-LA-EGFP or ONS-76-LA-EGFP cells transfected with siCTL or CD155 were implanted and then grown in the slices for 48 hours. For HGF stimulation, the feeding medium was supplemented with 20 ng/mL HGF. After treatment, the co-cultured slices were fixed with 4% paraformaldehyde and washed three times with PBS. Inserts were incubated in standard cell culture trypsin EDTA and incubated at 37 ^◦^C in a humidified incubator for 23 min. After 3 washes, the slices were blocked in PBS containing 3% fetal calf serum, 3% bovine serum albumin (BSA), and 0.3% Triton×100 for 1 hour at room temperature. Primary anti-Calbindin (Abcam, ab108404, 1:1000) and anti-human nuclei (Merck, MAB4383, 1:250) were diluted in the blocking solution and incubated overnight on a shaker at 4^◦^C. Proliferative cells were detected using the Click-iT EdU cell proliferation kit following (Invitrogen, C10340). Unbound primary antibody was removed with 3 washes with PBS supplemented with 3% BSA at RT. Secondary antibodies (Table S3) were incubated for 3 h at RT. The inserts were flat-mounted in glycergel mounting medium (Dako, C0563). Image acquisition was performed on an SP8 Leica confocal microscope (Leica Microsystems) and analyzed using Imaris (Oxford Instruments) and ImageJ (Fiji) software.

### Immunoblotting (IB)

Immunoblot was carried out as described in^14^. DAOY cells were prepared in 6 well plates to reach 70% confluency within 24 h. Standard media were replaced with SFM. After 24 h at 85% confluency, cells were treated with either SFM or SFM supplemented with 20 ng/ml HGF (PeproTech, 100-39) for 30 min. Cells lysis was performed with RIPA buffer for 15 min on ice and cleared lysates were analyzed by SDS-PAGE. Protein concentration was assessed using the Pierce BCA Protein Assay Kit (Thermo Fisher Scientific) according to the manufacturer’s instructions. Proteins were separated on Mini-Protean TGX (4-20%) SDS-PAGE gel (Bio-Rad) and transferred to PVDF membranes (Bio-Rad). Following 1 h of blocking with 5% non-fat milk, membranes were probed with primary antibodies listed in Supplementary material (Table S3). GAPDH was used as the internal loading control. HRP-linked secondary antibodies (1:2000) were used to detect the primary antibodies. Chemiluminescence detection was carried out using ChemiDoc Touch Gel imaging system (Bio-Rad). The integrated density of detected bands was quantified using Image Lab software (Bio-Rad).

### Boyden chamber assay

Transwells with 5 µm pore size (Sarstedt, 83.3932.500) were coated with 0.07 μg/μl Pure Col Collagen I (Advanced BioMatrix, 5005) dissolved in 70% EtOH for a final concentration of 10 μg.cm^-2^. 7’500 transfected cells (24 h after transfection) were resuspended in serum-free medium and seeded in the upper chamber of a 24 well plate containing the coated transwell. Medium in the lower chamber was either serum-free medium or serum-free medium supplemented with 20 ng/ml HGF (PeproTech, 100-39). Transwell migration was allowed for 18 hours, and cells were then fixed in 4% paraformaldehyde (Thermo Scientific, 28908) in PBS for 10 min. Fixed cells were stained with DAPI for 15 min. The remaining non-invading cells on the upper surface of the membrane were removed using a cotton swab. Images of DAPI-stained nuclei were acquired using an Axio Observer 2 plus fluorescence microscope (Zeiss) at 5x magnification. Nuclei were counted on the membrane areas using ImageJ software and plotted on Prism 9 software (GraphPad).

### Immunofluorescence analysis

1’500 transfected cells (24 h after transfection) were seeded in 384-well plate (Greiner Bio-One, 781090) and incubated overnight in normal growth conditions. Cells were starved in low serum condition (1% FBS) for 24 hours and treated with 20 ng/ml HGF for 10- or 30-min. Cells were fixed directly with 4% PFA at 37°C for 20 min. Between all subsequent steps, cells were washed five times with PBS using the 50 TS washer (BioTek). Cells were permeabilized with 0.05% saponin for 20 min at RT. Blocking was performed during 1 hour at RT with 1% FBS. Primary antibody (Table S3) solution was added to the sample and shaken at 100 rpm for 2 hours. Secondary antibody (Table S3) solutions were added to the sample and shaken at 100 rpm for 2 hours. Nucleic acids were stained with Hoechst-33342 (Sigma Aldrich, B2883, 1:2000 in PBS) for 20 min at RT. Image acquisition was performed using the Operetta CLS High-Content Analysis System (PerkinElmer, HH16000000) at 40x magnification. Nine fields per well with 9 z-planes with 500 nm z-spacing were acquired per site. Maximum intensity projection was computed and used for subsequent analysis. For puncta quantification, EEA1 or LAMP1 signals were segmented using the Harmony software (PerkinElmer) and quantified. The total number of puncta was referred to the number of cells present per field.

### Statistical analysis

The samples sizes and statistical tests were selected based on previous studies with similar methodologies. Sample sizes were not determined using statistical methods. Statistical analysis was performed using the Prism 9 software (GraphPad). One-way ANOVA repeated measures test using Tukey’s multiple comparison was performed for multiple comparisons. P-Value <0.05 were considered significant (* = p<0.05, ** = p<0.01, *** = p<0.001, **** = p<0.0001, ns = not significant).

## Data availability

The mass spectrometry proteomics data have been deposited to the ProteomeXchange Consortium via the PRIDE^72^ partner repository with the dataset identifier PXD030597.

## Author Contributions

C.C. and M.B. designed the study, prepared the figures, and wrote the manuscript (conceptualization, data curation, formal analysis, visualization, writing). C.C. planned the and conducted the majority of the experiments (investigation, methodology). J.M. generated the CRISPR-Cas9 MAP4K4 cells, conducted the SIA analysis with mitomycin C and generated samples and provided experimental support for *ex vivo* experiments (investigation, methodology). L.R. performed siRNA screening using the Boyden chamber assay (investigation, methodology). M.Z. contributed to the immunofluorescence, FACS and IB sample preparations and analyses (investigation, methodology). D.P. performed the patch clamping experiments (investigation, methodology, visualization) and S.N. contributed to the planning and evaluation of patch clamping experiments (methodology). A.G. generated samples and provided experimental support for *ex vivo* experiments and performed proliferation and viability analyses (investigation, methodology). MB was responsible for supervision and project administration, M.G. and M.B. managed funding acquisition.

## Acknowledgments

We thank Dr. Paolo Nanni, Dr. Jonas Grossman and Claudia Fortes from the Functional Genomic Center Zürich for their help and support for Mass Spectrometry analysis. We like to thank the UZH electrophysiology facility (e-phac) for their technical support. Imaging was performed with equipment maintained by the Center for Microscopy and Image Analysis, University of Zurich. This study was supported by grants from the Swiss National Science Foundation (SNF_31003A_165860/1, SNF_310030_188793) to MB and from the Childhood Cancer Foundation to MG.

## Supplementary files

**Table 1 | Surface Proteome Transmembrane prediction TMHMM filtering**

Excel File: Table 1_SurfaceProteome_TransMembraneAnnotation-TMHMM-Filtering_Capdeville2021

**Table 2 | Surface Proteome Plasma membrane annotation filtering**

Excel File: Table 2_SurfaceProteome_PlasmaMembraneAssociation_Capdeville2021

**Table 3 | Significant fold change for Transmembrane predicted proteins**

Excel File: Table 3_Significant_FoldChange_TransMembraneAnnotation_Capdeville2021

**Table 4 | Significant fold change for Plasma membrane annotated proteins**

Excel File: Table 4_Significant_FoldChange_PlasmaMembraneAssociation_Capdeville2021

**Tables S1-S3**

Antibodies and primer lists

---

**Figure S1.**
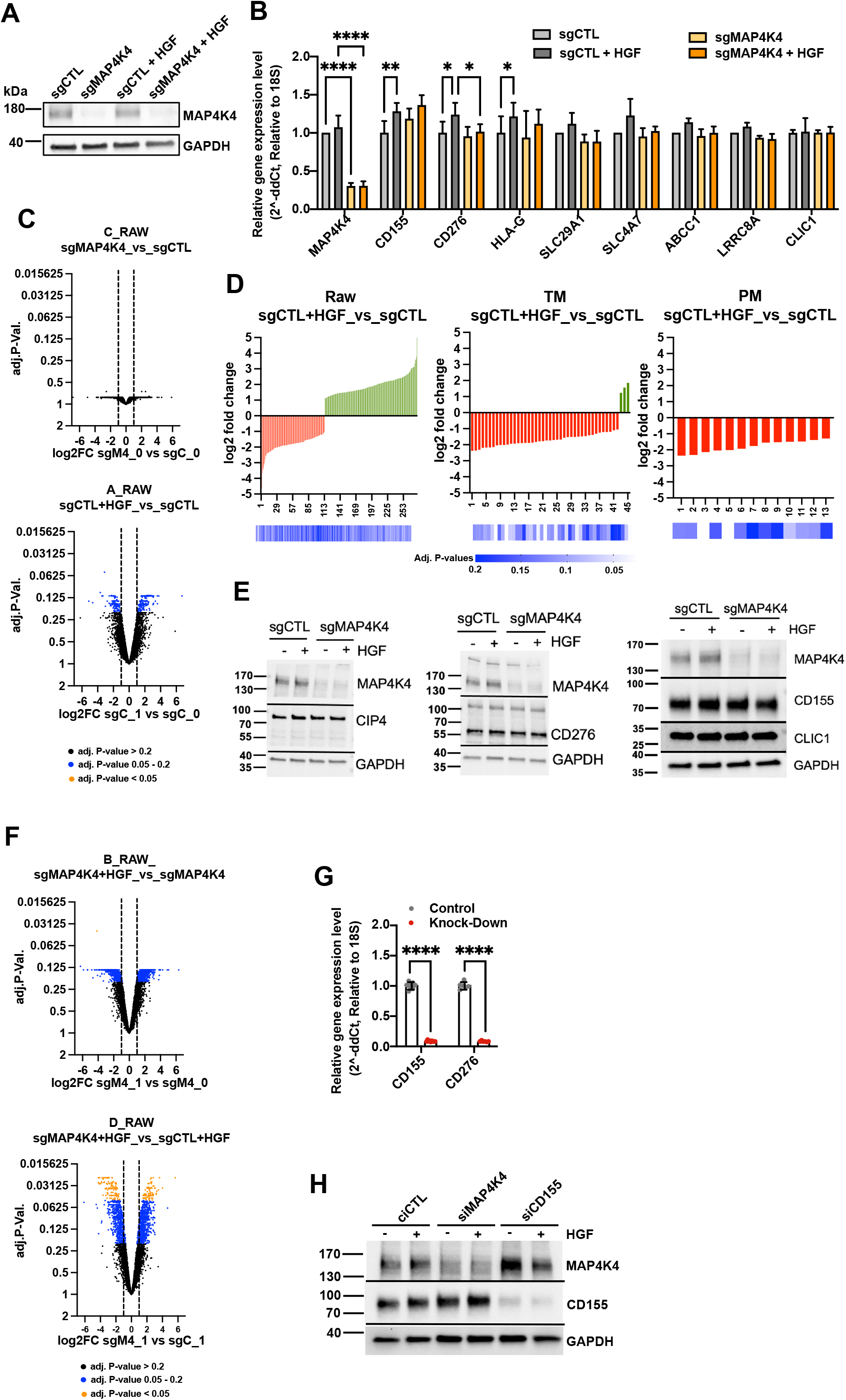
Validation of gene depletion efficiencies. **(A)** Immunoblot analysis of MAP4K4 expression in DAOY sgCTL and sgMAP4K4 cells used in proteome analysis. Treatments as indicated (HGF 20 ng/ml, 30 min). **(B)** Relative mRNA expression of a selection of proteins with significantly altered PM association. N = 3 independent experiments, mean & SD, * = p<0.05, ** = p<0.01, **** = p<0.0001. **(C)** Volcano plots of log2FC versus adjusted p-values of two-group analyses of all identified proteins. **(D)** Histogram of log2FC and adjusted p-values of two-group analyses. RAW: All proteins detected, TM: Transmembrane domain-containing proteins, PM: Plasma membrane-annotated proteins. **(E)** IB analysis of protein expression in sgCTL and sgMAP4K4 cells treated as indicated. **(F)** Volcano plots of log2FC versus adjusted p-values of two-group analyses of all identified proteins. **(G)** Relative mRNA expression analysis for *CD155* and *CD276* with siRNAs used. N = 3 independent experiments, mean & SD, * = p<0.05, ** = p<0.01, **** = p<0.0001 of unpaired t-test. **H)** IB analysis of CD155 expression in siMAP4K4 and siCD155 transfected cells.

**Figure S2.**
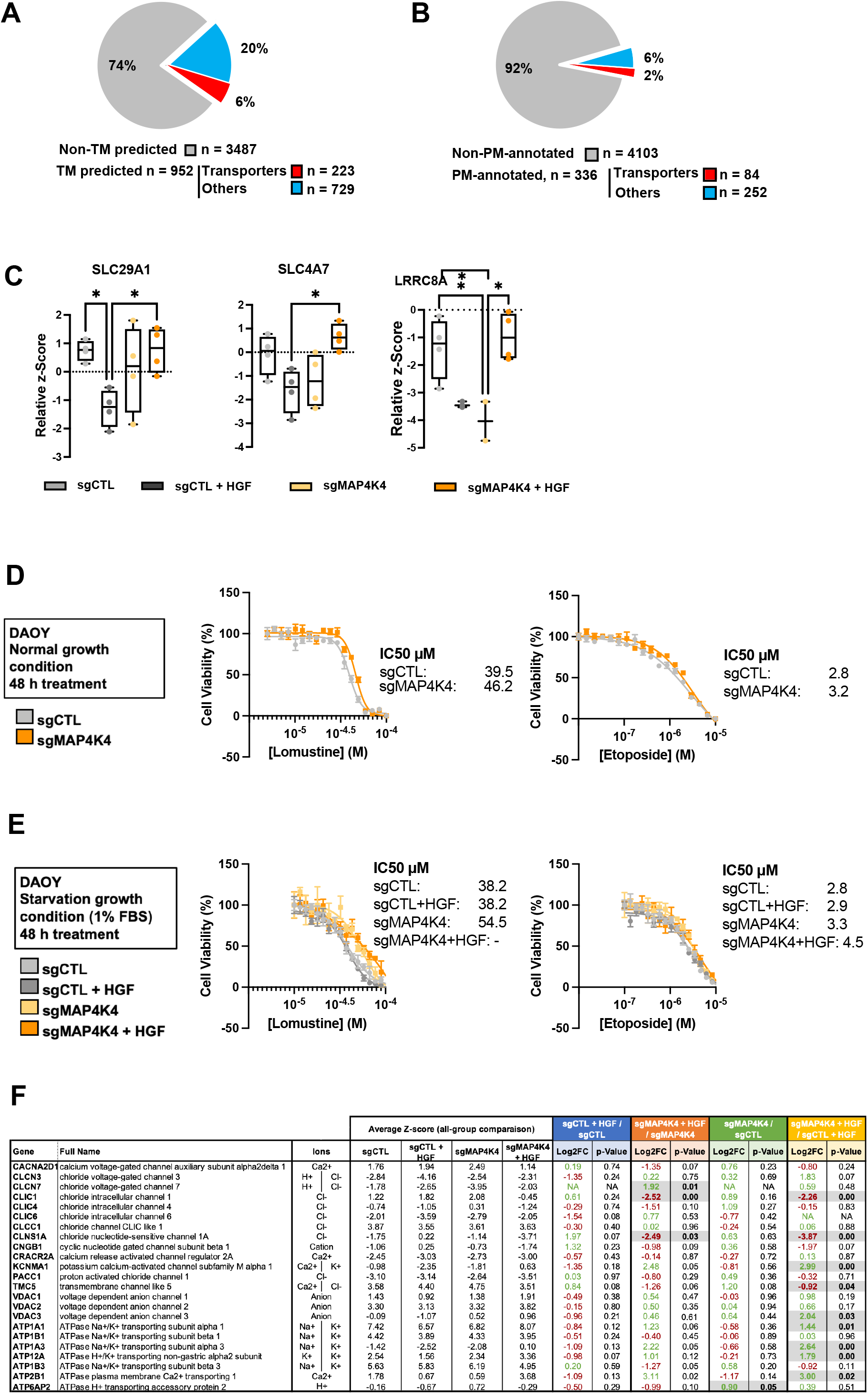
MAP4K4 promotes HGF-induced internalization of solute carriers and transporters. **(A)** Pie chart representing the proportion of proteins with predicted transmembrane domains (blue). Proteins annotated as transporters using Protein Atlas are highlight in red. **(B)** Pie chart representing the proportion of PM-annotaed proteins (blue) in all proteins identified in MS analysis. Proteins annotated as transporters using Protein Atlas are highlight in red. **(C)** Quantification of the PM-association of solute carriers and transporters involved in chemotherapy sensitivity and resistance. Relative z-Score value extracted from all-group comparison. n = 4 independent experiments, min to max, * = adjusted p<0.05 of one-way ANOVA. **(D)** Dose-response curve and corresponding IC_50_ values of Lomustine and Etoposide after 48 hours treatment under standard culture conditions (10% FBS). **(E)** Dose-response curve and corresponding IC_50_ values of Lomustine and Etoposide after 48 hours treatment +/- HGF stimulation under low-serum culture conditions (1% FBS). n ≥ 3 technical replicates, mean + SEM. **(F)** Selection of proteins with predicted transmembrane domains with knwon implication in ion transport. Potential ions involved are indicated. Normalized z-score values extracted from all-group comparison of MS analysis are depicted. Fold change and p-values extracted from each two-group analysis are shown. N = 4 independent experiments. Green: upregulated, red: downregulated, black: unchanged.

**Figure S3.**
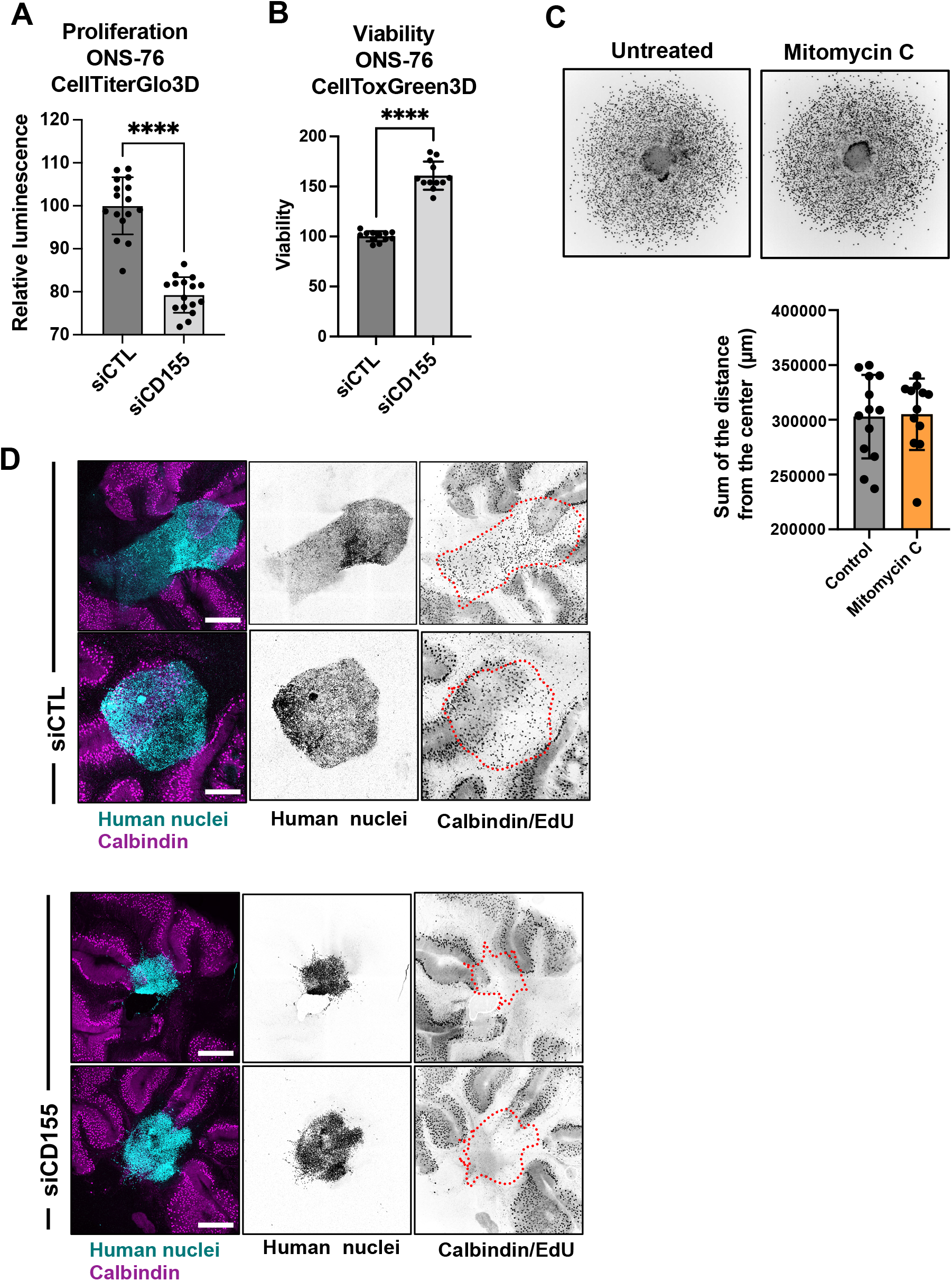
CD155 depletion prevents tissue invasion *ex vivo*. **(A)** CellTiter Glo proliferation assay of siCTL or siCD155 transfected ONS-76 cells grown as spheroids. **** = p<0.0001 of unpaired t-test. **(B)** CellTox Green cytotoxicity assay of siCTL or siCD155 transfected ONS-76 cells grown as spheroids. **** = p<0.0001 of unpaired t-test. **(C)** SIA analysis of ONS-76 cells treated with mitomycin C, 2 µM for 24h. Upper: Inverted grey scale images of Hoechst-stained nuclei at end-point. Lower: Quantification of total invasion total distance of invasion. **(D)** Confocal microscopy analysis of OCSCs two days after implantation of spheroid of DOAY cells transfected with indicated siRNAs. Cyan: Human nuclei, Magenta: Calbindin (Purkinje cells). Tumor cell area is indicated with dashed line in inverted gray-scale confocal images. Scale bar = 400 µm.

**Figure S4.**
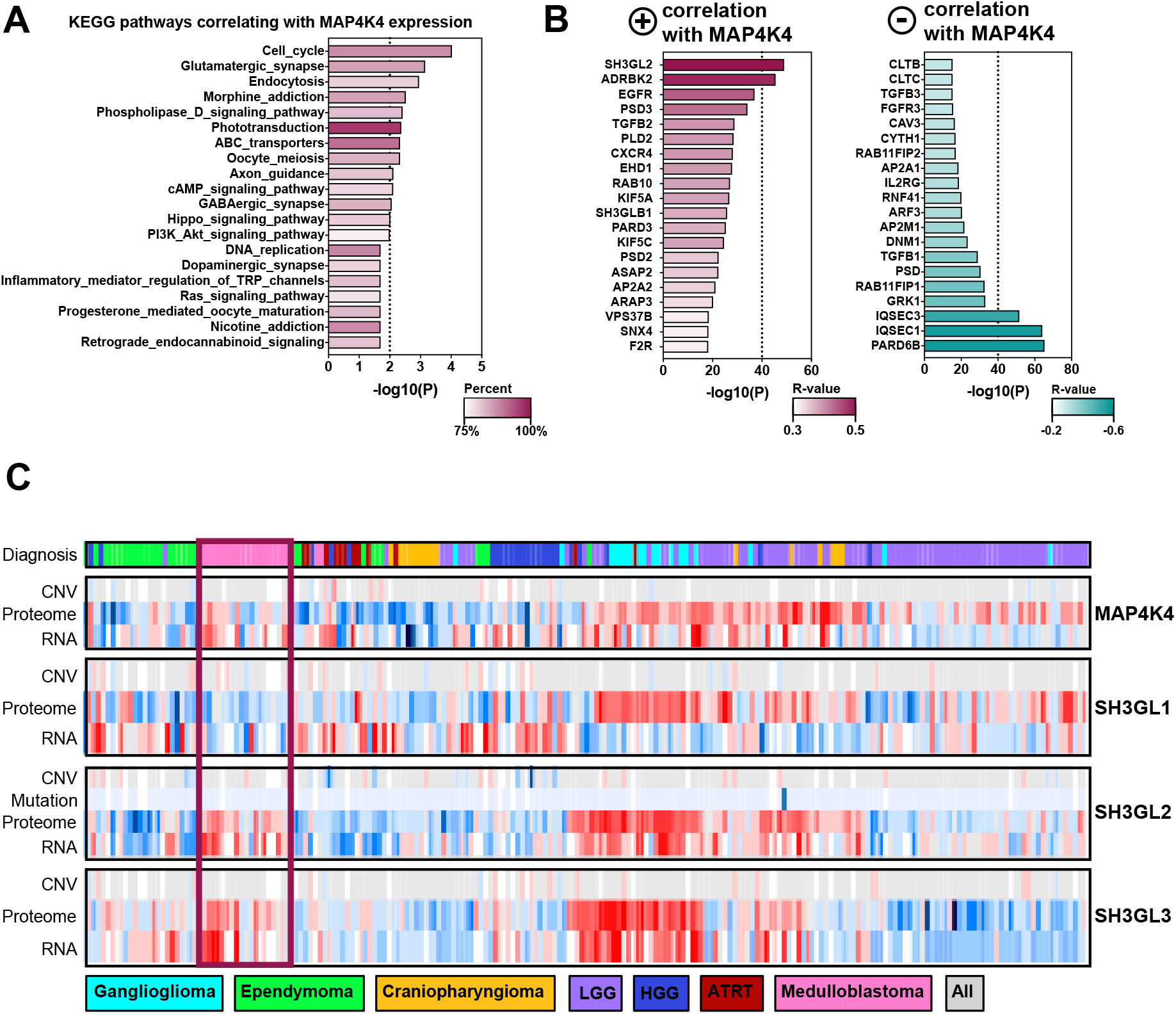
SH3GL2 is highly expressed in SHH MB tumors and correlates with MAP4K4 expression. **(A)** MAP4K4 co-expressed genes from 763 primary medulloblastoma samples were used to interrogate KEGG pathway database. **(B)** Top 20 genes with a positive (Left) or negative (Right) correlative expression score with MAP4K4. **(C)** Expression of MAP4K4, SH3GL1, SH3GL2 and SH3GL3 at RNA and protein levels in 218 pediatric brain tumors across seven histologic types. Copy number variation (CNV) and mutations are represented if available. The data were obtained from the R2: Genomics Analysis and Visualization platform (http://r2.amc.nl) and CPTAC data portal (http://pbt.cptac-data-view.org/).

**Figure S5.**
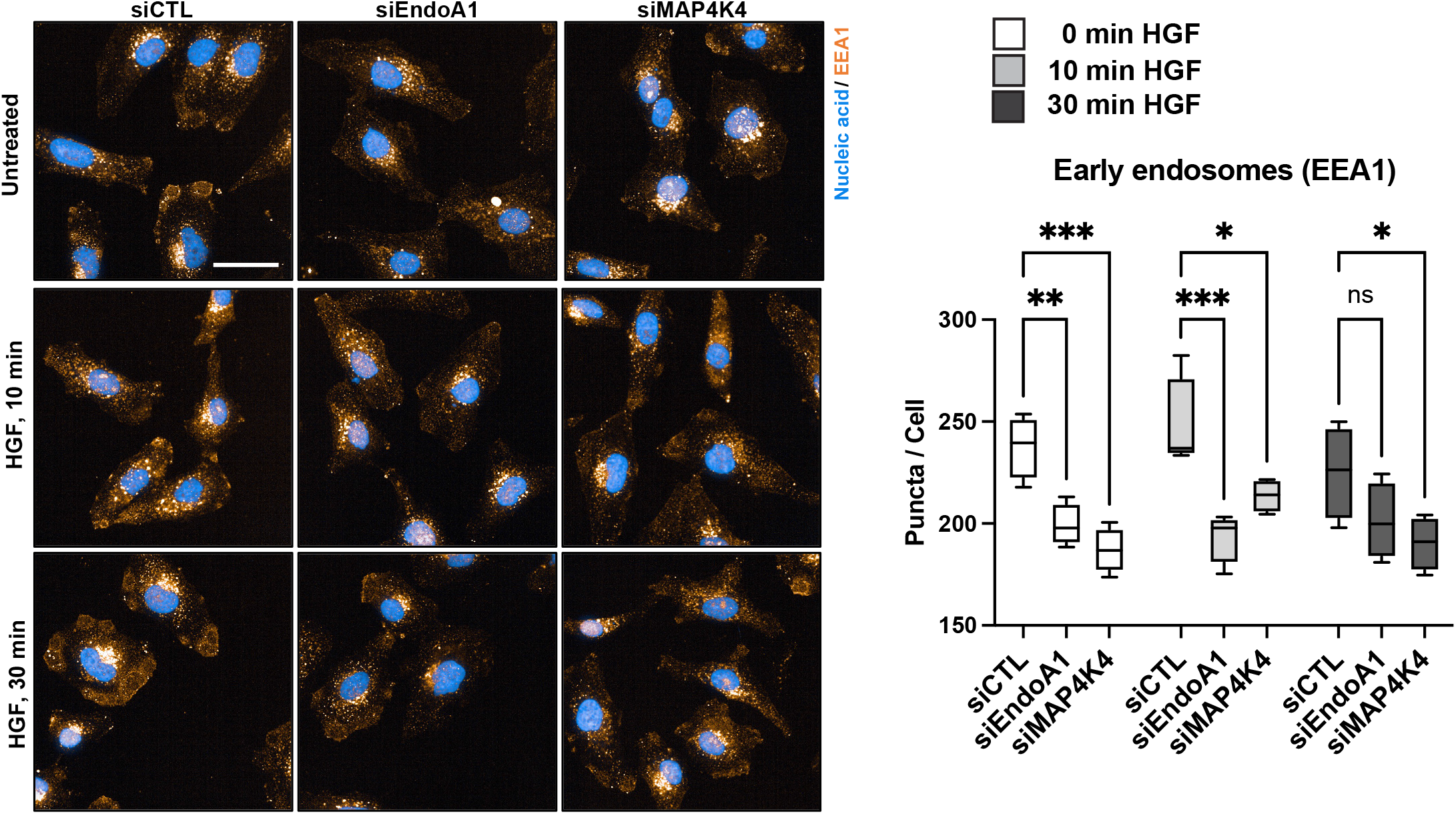
EndoA1- and MAP4K4-mediated regulation of EEA1. Confocal microscopy analysis of the early endosome marker EEA1. Left: Maximum intensity projection of siCTL, siEndoA1 and siMAP4K4 transfected DAOY cells. Blue: Nucleic acids (Hoechst), Orange: EEA1. Scale bar = 50 µm. Right: Quantification of the EEA1 positive puncta per cell. N = 4 technical replicates Min to Max, * p<0.05, ** p<0.01, *** p<0.001, **** p<0.0001, ns = not significant (one-way ANOVA).

**Figure S6.**
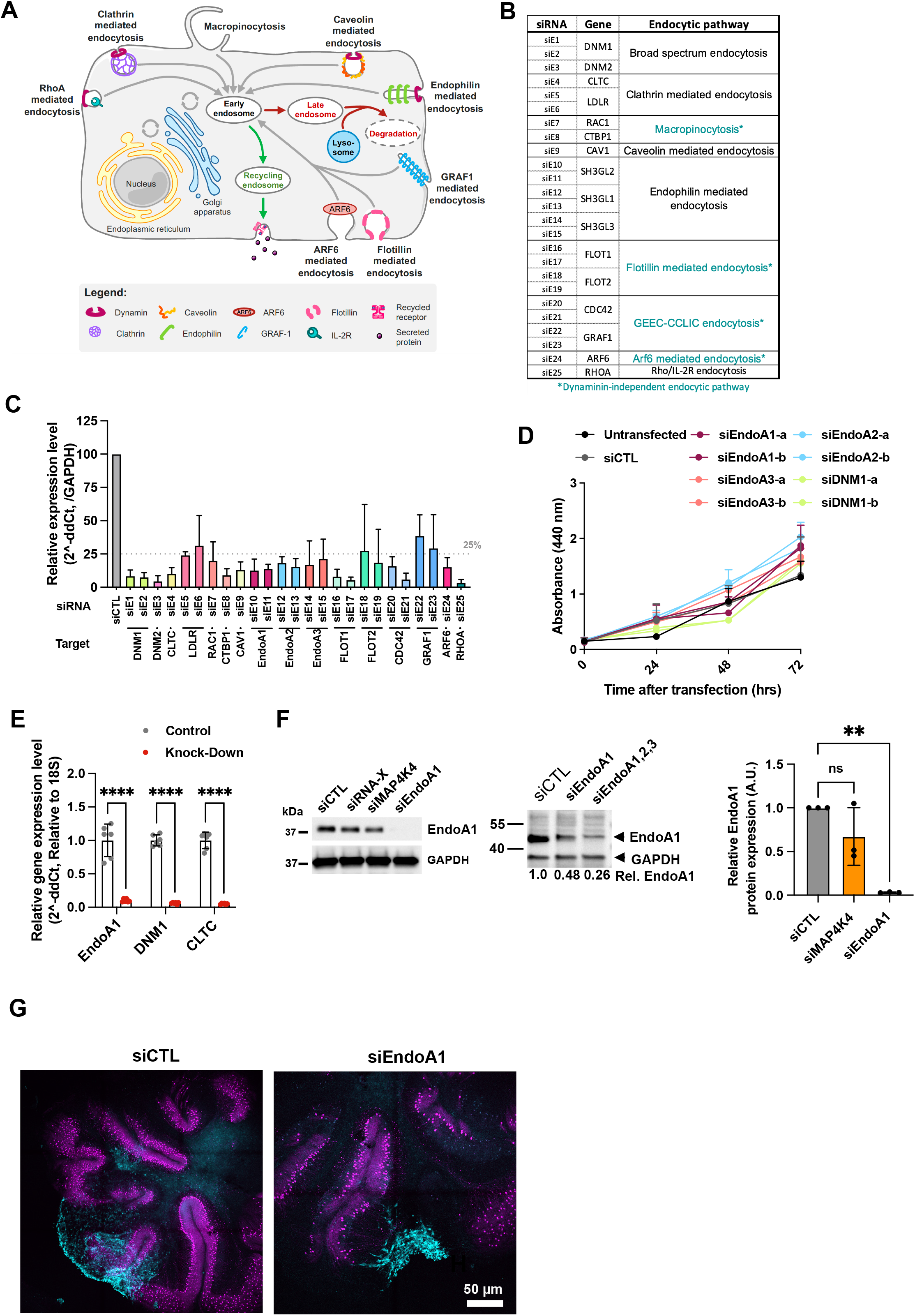
Implication of endocytosis on MB tumor progression. **(A)** Schematic representation of endocytic pathways in mammalian cells. **(B)** List of target genes used for siRNA screen and corresponding endocytic pathways. Pathways with dynamin-independent vesicle closure and abscission are shown in turquoise. **(C)** Depletion efficacy of siRNAs used in transwell migration screen assessed by RT-qPCR after transfection with the Dharmacon transfection reagent. RNA extraction occurred at endpoint of Boyden chamber assay. N = 2 to 9 independent experiments, Mean + SD. **(D)** WST cell proliferation analysis . N = 2 independent experiments, Mean + SEM. **(E)** Depletion efficacy of siRNAs against EndoA1, DNM1 and CLTC1 assessed by RT-qPCR with the Invitrogen transfection reagent. RNA extraction occurred at endpoint of SIA. N = 6 independent experiments, Mean + SD, **** p<0.0001. **(F)** Immunoblot (left and middle panels) and RT-qPCR analysis (right panel) of siRNA-mediated Endophilin-A1 depletion in ONS-76 cells. Quantification of RT-qPCR analysis: N = 3 independent experiments, Mean + SD. ** p<0.01, ns = not significant. **(G)** Confocal microscopy analysis of OCSCs two days after implantation of spheroid of DOAY cells transfected with indicated siRNAs. Cyan: LA-EGFP, Magenta: Calbindin (Purkinje cells).

